# Climate benefits of seaweed farming: estimating regional carbon emission and sequestration pathways

**DOI:** 10.1101/2023.06.13.544854

**Authors:** Cameron D. Bullen, John Driscoll, Jenn Burt, Tiffany Stephens, Margot Hessing-Lewis, Edward J. Gregr

## Abstract

Seaweed farming is widely promoted as an approach to mitigating climate change despite limited data on carbon removal pathways and uncertainty around benefits and risks at operational scales. We explored the feasibility of seaweed farms to contribute to atmospheric CO_2_ reduction in coastal British Columbia, Canada, a region identified as highly suitable for seaweed farming. Using a place-based, quantitative model, we examined five scenarios spanning a range of industry development. Our intermediate growth scenario sequestered or avoided 0.20 Tg CO_2_e / year, while our most ambitious scenario (with more cultivation and higher production rates) yielded a reduction of 8.2 Tg CO_2_e /year, equivalent to 0.3% and 13% of annual greenhouse gas emissions in BC, respectively. Across all scenarios, climate benefits depended on seaweed-based products replacing more emissions-intensive products. Marine sequestration was relatively inefficient in comparison, although production rates and avoided emissions are key uncertainties prioritized for future research. Our results show how seaweed farming could contribute to Canada’s climate goals, and our model illustrates how farmers, regulators, and researchers could accurately quantify the climate benefits of seaweed farming in local contexts.

## Introduction

Climate change, driven largely by increasing atmospheric carbon dioxide, is now one of the greatest challenges threatening humanity and global ecosystems ^[1,2]^. Carbon dioxide removal (CDR) strategies are increasingly seen as necessary for meeting global climate targets ^[3]^, with seaweed aquaculture recently gaining attention as a promising approach ^[4–6]^. This interest in seaweed is due to the high productivity of many species and their efficiency at drawing CO_2_ from the water and converting it into organic biomass ^[7]^. Oceans also play a significant, natural role as a carbon sink, taking up an estimated 2.8 Gt C / year in the 2011-2020 period, equivalent to approximately 30% of annual fossil fuel emissions ^[8]^. As such, several strategies have emerged to try to enhance the rate of carbon sequestration and storage in the ocean by protecting, restoring, or enhancing productivity of wild marine plants, macroalgae, and phytoplankton ^[6,9,10]^.

Recently, large-scale farming of seaweed has been put forward as a potential CDR strategy, with various groups including the International Panel on Climate Change highlighting seaweed aquaculture as an important area for research and development ^[4,11–13]^. In response, a variety of approaches to farming seaweed for the express purpose of CDR have been proposed, including the purposeful transport of seaweed biomass to the deep ocean where it can remain for long periods ^[14–17]^, and the use of seaweed to produce lower emission products such as biofuel ^[18,19]^. Such approaches remain largely untested, and their efficacy and ecological impacts remain uncertain ^[6,20,21]^.

There are various pathways by which seaweeds, whether wild or farmed, may sequester carbon in the marine environment ^[20,22]^. Much of the net primary production of wild seaweed, over 80% by some estimates ^[23]^, is released as detrital particulate and dissolved organic carbon (POC and DOC) ^[24,25]^. Much of this POC and DOC is consumed and recycled, nourishing coastal ecosystems and driving secondary production ^[26,27]^. Unconsumed, recalcitrant POC and DOC can be transported to nearby sediments or exported by currents into deeper water where it is sequestered ^[22,28,29]^. Unlike wild seaweeds, farmed seaweed is typically harvested after the growing season, likely reducing the POC and DOC produced ^[30]^. Recent proposals to actively sink seaweed, either around farms or in deep waters, aim to facilitate and enhance this natural sequestration process ^[5,31]^.

Beyond ocean dumping, harvested seaweed biomass would help reduce atmospheric greenhouse gas emissions if used as a replacement for products that use more land, water, and carbon resources. Such products include foods and food additives, animal feed, biofuels, soil additives (biochar and biostimulants), and pharmaceuticals and cosmetics ^[32–34]^. Seaweed aquaculture typically requires minimal inputs of material and energy, uses no fertilizer, and produces limited emissions and can therefore help decarbonize production systems by replacing carbon-intensive alternatives ^[35–37]^. A large proportion of global seaweed production, approximately 31–38% ^[38]^, is currently used for human consumption and there is substantial interest in using seaweed to create more sustainable food systems and supply chains ^[36,37]^. Other uses of seaweed such as cement additives or biochar for agriculture ^[34]^ can directly sequester carbon in built environments and soils. However, the ability for seaweed products to reduce emissions by replacing higher intensity products depends critically on the emissions profile of the food or product that seaweed is replacing, the efficiency of seaweed production and processing, and a sufficient market for the seaweed products.

Given these various pathways for climate benefit from seaweed aquaculture, there is mounting interest in the potential climate benefits this industry might offer. Seaweed aquaculture currently accounts for 29% of the global aquaculture production by weight (FAO 2022) with the vast majority (over 99%) currently grown in Asia ^[21,39]^. Recent estimates have suggested that between 48–119 million km^2^ of the global ocean (an area 24–60 times the size of Greenland) may be suitable for seaweed production ^[5,40]^, however the industry remains nascent in most countries ^[5,41]^. Farming seaweeds also has other potential benefits including reducing excess anthropogenic nutrients in ocean water, nourishing coastal ecosystems, and reducing wave impacts on shorelines ^[4,42,43]^. As such, there is considerable excitement around seaweed aquaculture and substantial marketing, investment, and media attention focused on the climate and ecosystem benefits of the industry ^[16,44]^. Given this enthusiasm, there is an urgent need to ensure our understanding of the benefits as well as risks of seaweed aquaculture informs potential industry development ^[21,45]^.

In this study, we begin to address several knowledge gaps around the efficacy of sequestration pathways and the associated emissions by developing a place-based, data-driven, mathematical model to explore the potential of seaweed aquaculture to reduce atmospheric greenhouse gases. We consider several CO_2_ sequestration and emissions pathways with a focus on production and processing emissions, and marine sequestration. We applied our model to a case study in Canada, a nation identified to have extensive – but unexplored – potential to develop seaweed aquaculture for climate mitigation purposes ^[41,46]^.

We parameterized our model and developed scenarios based on a case study of kelp (seaweeds of order Laminariales) aquaculture in British Columbia (BC) - a province with a long coastline of nutrient rich waters, abundant wild seaweeds suitable for cultivation, and a growing kelp farming industry. Using a suite of increasingly ambitious scenarios representing increasing degrees of aquaculture development and technological advances to estimate a range of net annual atmospheric draw-down values (Tg CO_2_e / year). We grounded our model in discussions with regional kelp producers, using parameter values obtained from current kelp aquaculture operations and the published literature (See Methods and Supplementary Material).

## Results

### Aquaculture Development Scenarios

We used five scenarios to reflect a range of kelp aquaculture development in BC (Table 1). For each scenario we calculated the spatial extent of potential kelp farms using assumptions about suitability (depth and substrate), access to coastal communities and infrastructure, and overlap with other human uses (Fig. 1). Spatial extents ranged from 507 km^2^ in modest development scenarios to 5,681 km^2^ in our most extensive scenario. We used a combination of survey responses and unstructured interviews with kelp producers in BC and Alaska to ground these scenarios in species profiles, farm footprints, production rates, emission sources, and seaweed fates common to the temperate eastern North Pacific. All scenarios use a species mix (80% *Saccharina latissima*, 10% *Alaria marginata*, and 10% *Nereocystis luetkeana*) in line with current BC production. The scenarios varied in the spatial extent of kelp aquaculture, kelp production rates, and the fate of kelp biomass (Table 1).

**Table 1.**
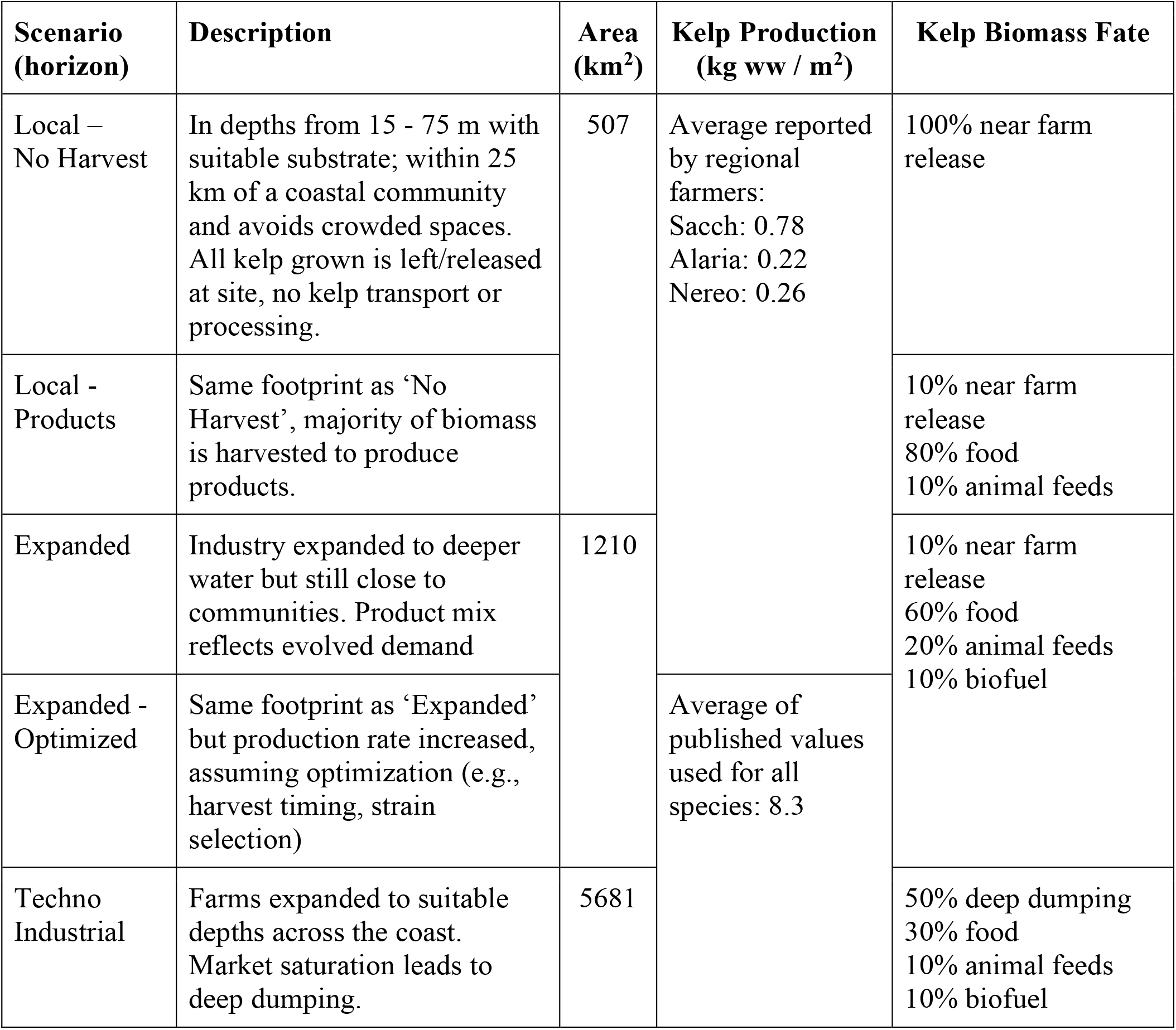
Summary of the five scenarios examined. For each scenario, the size of the growing area, the mean rate of kelp production, and the fate of kelp biomass is outlined.

**Fig. 1.**
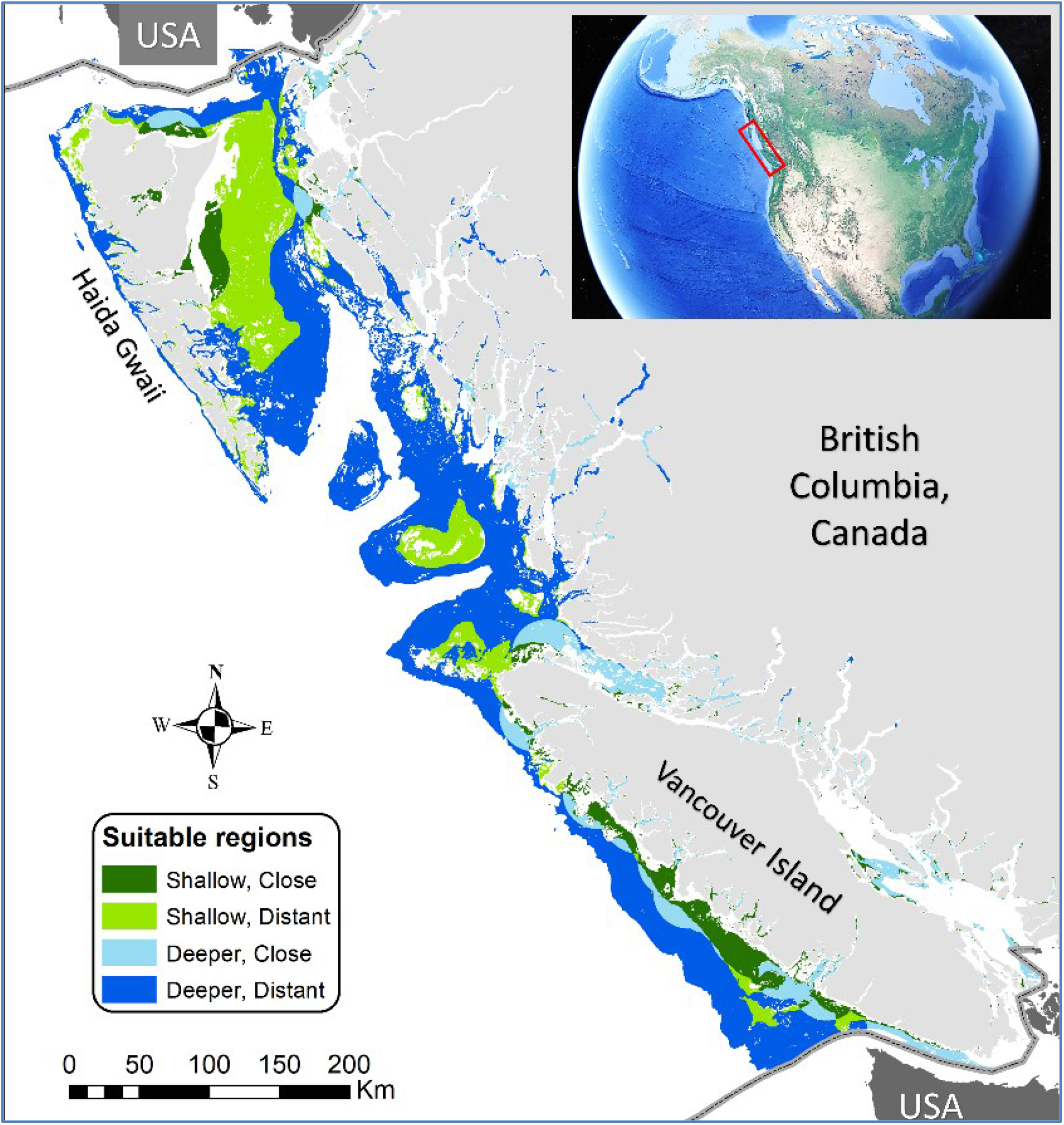
Potentially suitable areas for seaweed aquaculture in BC, Canada. Our study area showing suitable locations cultivated under different production scenarios (main panel) and the location of our study in North America (inset). We defined suitable areas as those with soft substrates, at optimal depths (≥ 15 and ≤ 200 m), with low human use. Shallow suitable waters close to communities (dark green) are assumed to be favored under the Local scenarios; the areas available for cultivation under the Expanded scenarios also include deeper waters close to communities (light blue). The Techno Industrial scenario includes cultivation across all suitable areas. Small pockets of shallow areas close to communities on the eastern side of Vancouver Island, as well as deeper locations in the mainland inlets are notable for early kelp farm development. See text for details on the scenarios and our methods for identifying suitable areas.

### Net Climate Benefits

Our simulations indicate that kelp aquaculture can provide substantial climate benefits, although there is large variability in model estimates. We present our results for each model scenario as the median value from 10,000 Monte Carlo runs, along with the 25^th^ and 75^th^ percentiles. As an example of an intermediate level of industry development, the *Expanded* scenario (Fig. 2) estimates a net atmospheric reduction of 0.196 (0.084-0.345) Tg CO_2_e / year (results for other scenarios are provided in Table S1). This climate benefit is achieved by producing 0.969 (0.572-1.45) Tg ww of harvestable kelp per year, directing 80% of it to seaweed-based products (primarily food and animal feed), and leaving 20% of it in the water where it may be consumed, re-mineralized, or eventually sequestered. In this scenario, seaweed products avoid the release of 0.29 (0.17-0.44) Tg CO_2_e / year from the replacement of existing carbon-intensive products, while the biomass left in the water sequesters only 0.0012 (0.0007-0.002) Tg CO_2_e / year, about 200 times less. From an estimated loss of 0.023 (0.012-0.041) Tg of POC and DOC during production, we estimate about half – 0.011 (0.0057-0.020) Tg CO_2_e / year – would be sequestered. We predicted nursery operations would emit 0.011 (0.007-0.014) Tg CO_2_e / year; at-sea cultivation operations to emit 0.060 (0.050-0.070) Tg CO_2_e / year; and transport and processing of harvested seaweed to emit 0.032 (0.017-0.053) Tg CO_2_e / year.

**Fig. 2.**
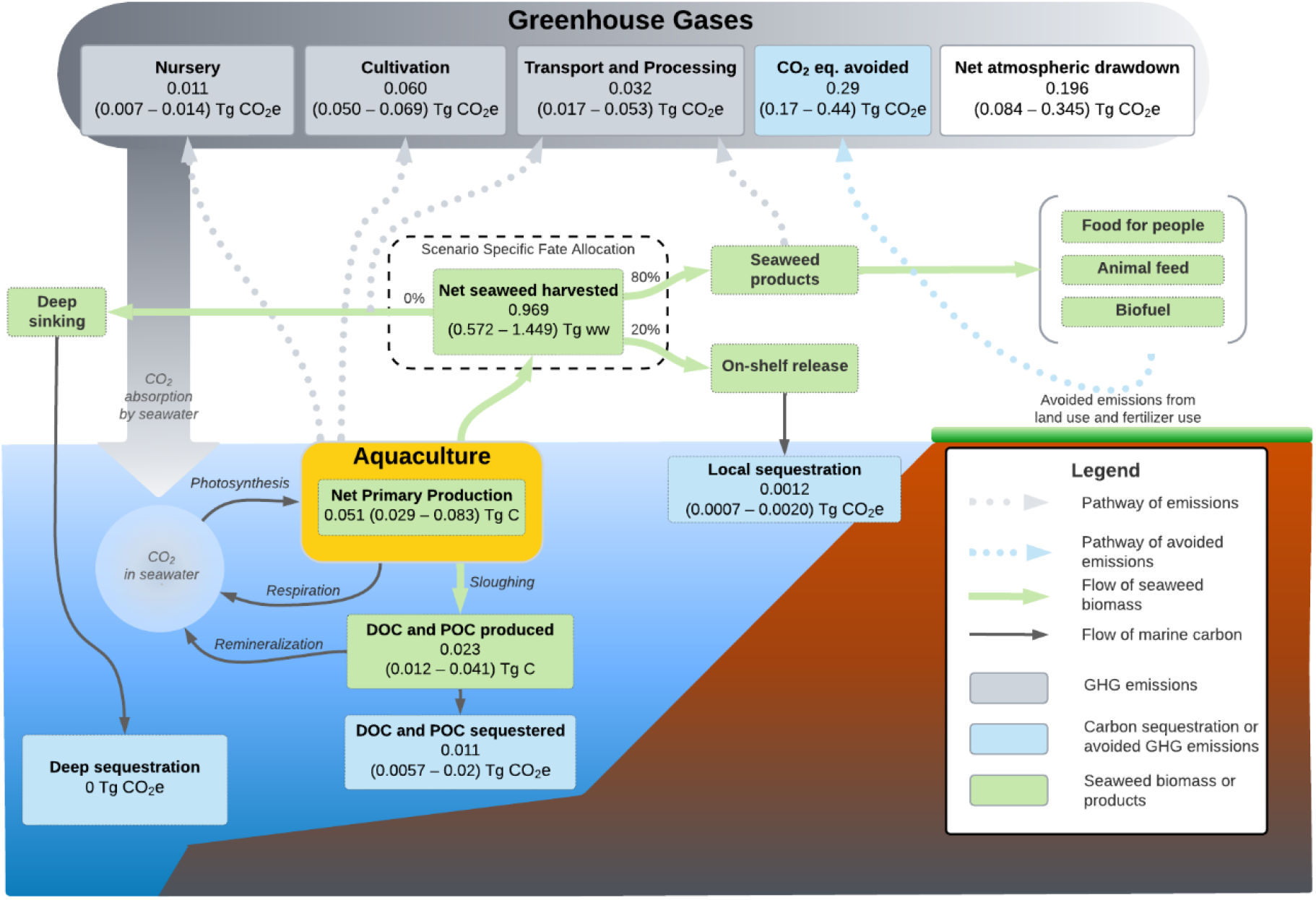
Illustrative model structure. Diagram illustrating the structure and carbon pathways represented in the mathematical model. Results for key model elements are shown for the Expanded scenario as median (25th percentile - 75th percentile) estimates from 10,000 Monte Carlo runs. Results for all scenarios are provided in Table S1. Processing emissions not considered include product packaging, storage, and transport and other unquantified life cycle components (see supplementary material).

### Differences between scenarios

Model estimates varied by scenario due to differences in spatial extent, production rates, and the fate of harvested kelp (Fig. 3). Under our most conservative scenario (*Local – No Harvest*) kelp aquaculture was a net emitter of CO_2_, producing 0.02 (0.010-0.023) Tg CO_2_e / year and suggesting total emissions from production would exceed this scenario’s sequestration potential (Supplementary Fig. S1 and S2, Table S1). In contrast, our most developed scenario (*Techno Industrial*) resulted in a net draw down of 8.15 (5.39 -11.53) Tg CO_2_e / year. This was primarily comprised of emissions avoided by seaweed-based products (8.9 [6.0-12.3] Tg CO_2_e / year) and to a lesser extent biomass actively sequestered in deep water (0.71 [0.44-1.10] Tg CO_2_e / year (Supplementary Fig. S1 and Table S1). Total emissions for this scenario were estimated at 1.40 (0.99-1.92) Tg CO_2_e / year.

**Fig. 3.**
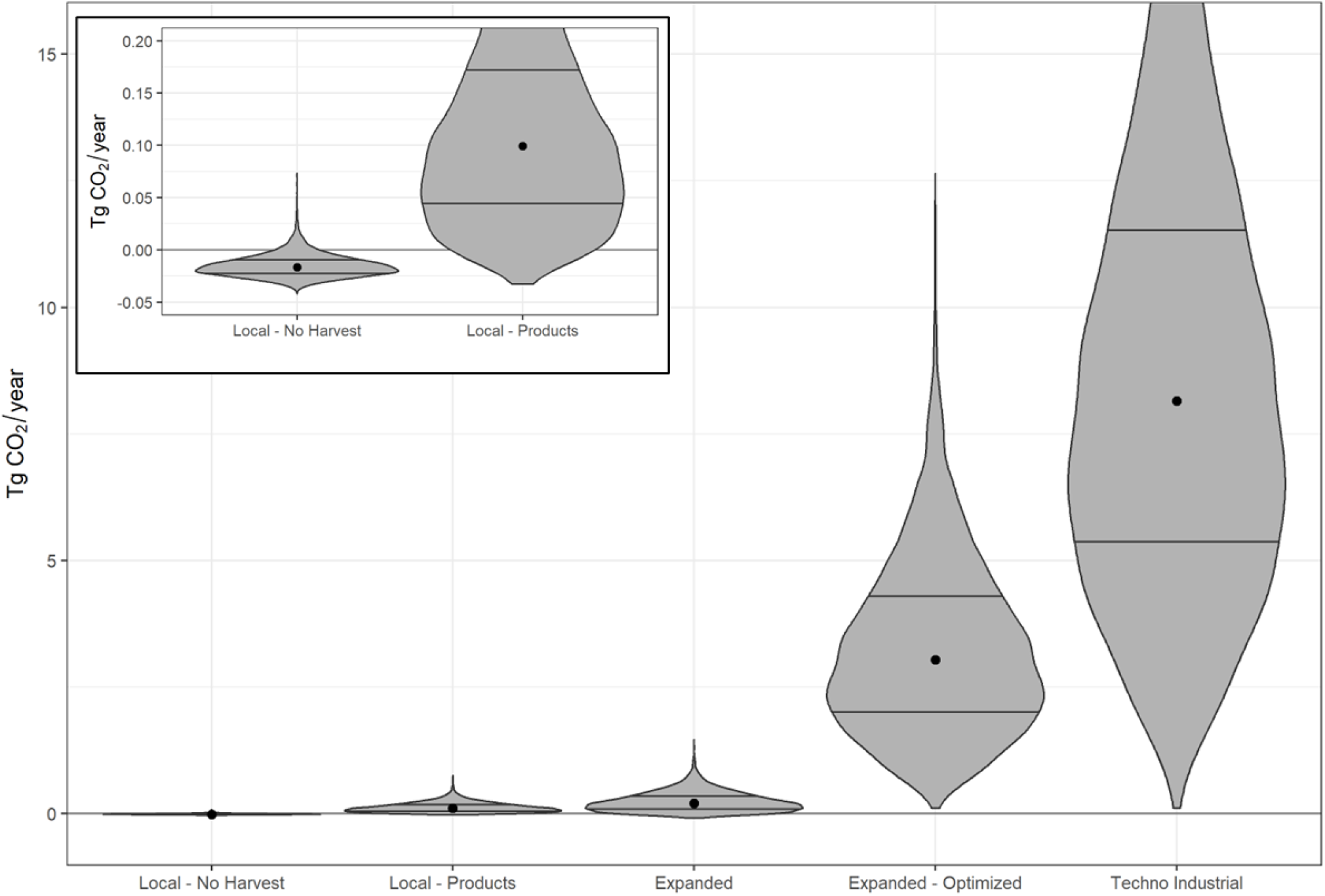
Net reduction in atmospheric CO_2_ for each scenario. Violin plots illustrate the distribution of estimates from 10,000 Monte Carlo runs, with the central dot indicating the median and horizontal lines at the 25^th^ and 75^th^ percentiles. Inset plot shows a re-scaled version of the two Local scenario results.

The *Local – Products* and *Expanded – Optimized* scenarios fall between these two extremes and are closer to the *Expanded* scenario. The *Local – Products* scenario estimates a net reduction of 0.10 (0.045 - 0.172) Tg CO_2_e / year while the *Expanded – Optimized* scenario yields a net reduction of 3.04 (2.02 to 4.29) Tg CO_2_e / year. Uncertainties are high and the range of plausible values is large for all model scenarios.

### Sensitivity Analysis

We applied two sensitivity analyses to examine uncertainty in the model. We first assessed the influence of uncertainty at the scale of the three main sub-models (production, emissions, and products-sequestration). The second analysis examined the sensitivity of model results to uncertainty in the individual model parameters within each sub-model. These results are scenario-specific as they depend on the scenario configurations (results for all scenarios are provided in Supplementary Fig. S3-S4).

Of the three sub-models, uncertainty in production contributed most to the overall model uncertainty (Fig. 4A; *Expanded* scenario). Uncertainty in sequestration parameters contributed a smaller amount while emission parameters contribute the least to overall model uncertainty. The parameter-level sensitivity analysis identified the parameters with the highest uncertainty in each sub-model (Fig. 4B). Species-specific production rates contribute the most uncertainty to the production sub-model, most obviously with Saccharina as it’s the dominant species farmed. In some iterations, variability in the production rate of Saccharina can result in more than a 4-fold increase in the estimated net CO_2_ reduction (Fig. 4B). In the sequestration sub-model the food emission replacement factor had the highest uncertainty, with estimates of net CO_2_ reduction varying by more than ±50% depending on the value of this parameter. Parameters estimating material production (concrete, steel, etc.), energy use, and seaweed processing contributed the most uncertainty to the emissions sub-model. However, this sub-model contributed little to the overall uncertainty in our model.

**Fig. 4.**
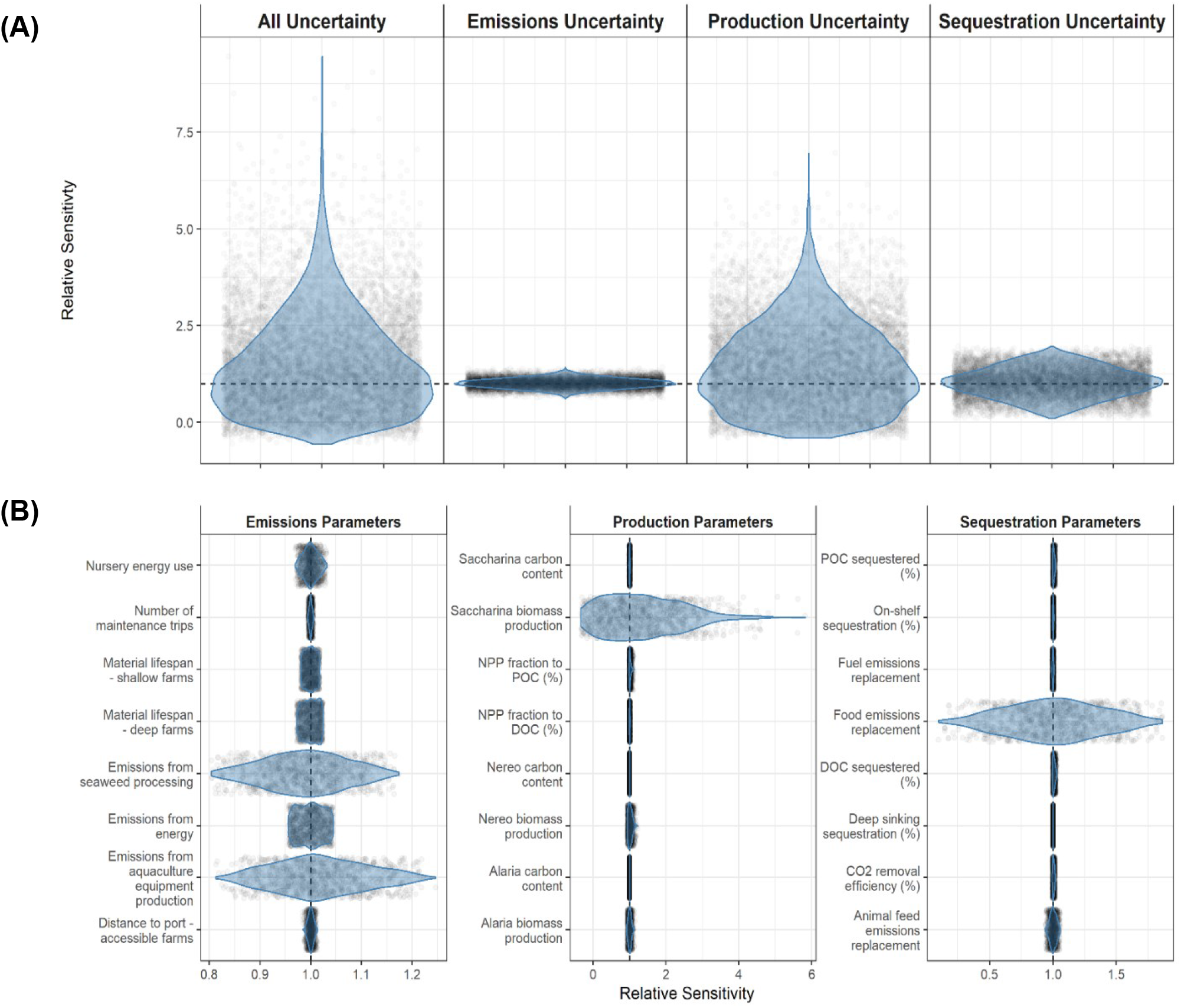
Relative sensitivity of the estimated net reduction in atmospheric CO_2_. Results are shown for the *Expanded* scenario, as (**A**) cumulative uncertainty in different parameter categories, and (**B**) uncertainty in individual parameters. Sensitivity is assessed relative to a model run with no uncertainty, where model parameters were set to their central estimate, yielding a net CO2 reduction of 0.155 Tg CO_2_ / year. Violin plots show the distribution of the Monte Carlo runs, with each individual estimate shown as a shaded point. Note the plots in panel **B** have different x-axis scales.

## Discussion

We used our model to explore a range of seaweed aquaculture development scenarios and various pathways for sequestering carbon or avoiding CO_2_ emissions. Our findings indicate that, while not a silver bullet, seaweed aquaculture may have a role in helping to reduce greenhouse gas concentrations, particularly if biomass is used to create products such as food that can replace traditional, high emission products. By grounding the model in the local context of BC, we also provide, for the first time, an estimate of the potential climate benefits from kelp aquaculture in Pacific Canada. Our sensitivity analyses highlight where targeted future research would improve our understanding of seaweed aquaculture systems.

A key finding from our model is that seaweed-based products have substantially more potential for reducing atmospheric greenhouse gas emissions through product replacement than passively or actively sinking seaweed for carbon sequestration. Given high rates of re-mineralization, inefficient transport to depth, and current uncertainties around how much of the carbon in seaweed biomass is measurably responsible for atmospheric drawdown ^[20,47]^, we show that marine sequestration pathways are likely to be inefficient with only a small fraction of seaweed biomass sequestered for the long term (> 100 years). In comparison, seaweed-based products can replace existing products that have high, well-described emissions such as agricultural land use and nutrient management ^[48]^ resulting in significant reductions in atmospheric greenhouse gases. On a marginal (per-area) basis, we estimate that with optimized production seaweed farmed to replace traditional food products (e.g., oil crops, pulses, or cereals) could avoid 3,238 (2085 - 4696) T CO_2_e / km^2^ / year while sinking seaweed biomass in deep water would sequester only 296 (164 - 483) T CO_2_e / km^2^ / year (Supplementary Table S2). These estimates show good concordance with values from the literature: recent work by Spillias et al. ^[40]^ found a similar magnitude of potential benefits for food replacement pathways, while sequestration in deep water has previously been estimated to sequester between 60.3 T CO_2_e / km^2^ / year ^[31]^ and 1,110 T CO_2_e / km^2^ / year ^[5]^.

Our *Local – No Harvest* scenario emphasizes this result as it suggests that leaving all farmed kelp biomass in the water may generate more CO_2_ than would be sequestered. Actively sinking kelp in deep water (e.g., the *Techno Industrial* scenario) results in almost four times more sequestration but remains much less effective than the production of kelp-based replacement products (Supplementary Table S2). Thus, our results strongly suggest marine carbon sequestration alone is unlikely to justify kelp aquaculture as a CDR strategy. Ethical concerns have also been raised around purposeful sinking of seaweed, as this nutritionally valuable biomass could play a role in improving regional and global food security ^[45]^.

The climate benefits of seaweed-based products are largely dependent on their ability to replace products with higher associated emissions, including the existence of a sufficient market.

Although we assumed a robust market for kelp-based products, the costs of kelp production, the market price, and any related incentives (e.g., carbon credits) will play a key role in determining the development of this industry. Further, while we were able to include several major sources of emissions, we could not explicitly consider downstream emissions from seaweed-based products such as product packaging, product transport, and waste management as these emissions require detailed life cycle analyses and depend heavily on local context. Similarly, the values used to parameterise the emissions offset by seaweed replacement products are largely limited to emission reductions due to land use and nutrient management. Additional associated emissions (e.g., those related to fertilizer production, agricultural energy use, and product storage and transport) would also be avoided when replacing traditional products but are difficult to estimate. Considering a broader range of emissions related to both seaweed and traditional (potentially replaceable) products will provide greater insight into the climate benefits of seaweed-based products. To ensure a conservative assessment of the climate benefits from seaweed aquaculture, we did not consider more nascent products, such as methane-reducing cattle feed additives which, while promising ^[40,49]^, remain uncertain and difficult to quantify.

With an appropriate mix of seaweed fates, our results show that modest development scenarios (*Local – Products* and *Expanded*) reduce atmospheric greenhouse gases by 0.10 and 0.20 Tg CO_2_e / year, respectively. These reductions correspond to 0.15 – 0.3% of total annual emissions from BC (64.6 Tg CO_2_e / year in 2020 ^[50,51]^). Our most ambitious scenario (*Techno Industrial*) led to a reduction of 8.2 Tg CO_2_e / year, or 12.6% of BC’s annual emissions. This substantial range of possibilities for kelp aquaculture shows that reducing or offsetting society-wide emissions with kelp aquaculture must be weighed against other costs and benefits (see below).

Our predicted estimates of CO_2_ reduction by kelp aquaculture in BC compare favourably to recent estimates for natural climate solutions across Canada ^[52]^. Specifically, estimates from our *Local - Products* scenario are similar to strategies such as seagrass restoration (0.1 Tg CO_2_e/year in 2050), while our maximalist *Techno Industrial* scenario compares to more substantial strategies such as improved agricultural nutrient management or the use of cover crops (6.3 and 9.8 Tg CO_2_e/year in 2050, respectively)^[52]^. On a national scale, considering all three of Canada’s oceans (including the Atlantic and Arctic) would substantially increase the potential climate benefits of kelp aquaculture. An additional consideration is that unlike many terrestrial natural climate solutions, kelp aquaculture avoids potential conflicts between other competing land uses such as food and bioenergy production ^[52,53]^ although it will compete for space in the marine environment ^[54]^. Kelp aquaculture may even facilitate terrestrial natural climate solutions by alleviating reliance on terrestrial agriculture – making space for land based conservation, restoration, and regenerative land management ^[40]^.

The scenarios we explored used between 507 km^2^ to 5,681 km^2^ of BC’s coastal waters for kelp aquaculture, areas approximately equivalent to the Isle of Man, UK and Prince Edward Island, Canada, respectively. While these extents represent substantial growth of the aquaculture industry from present day, they are conservative as our estimate of total suitable area in BC is as much as 10 times that (Fig. 1) and other recent estimates suggest widespread suitability ^[40]^.

Potential climate benefits could thus be much higher than our estimates, but development to the full suitable extents would come with substantial social and ecological risks.

While our goal was to assess the potential climate benefits of kelp aquaculture in BC, its feasibility will also need to consider a number of societal, economic, and ecological questions. Seaweed aquaculture has often been shown to provide sustainable economic livelihoods and contribute to community well-being, however this is dependent on local and regional conditions ^[42,55,56]^. In BC, seaweed aquaculture expansion would likely require development in remote coastal communities (e.g., transport costs may incentivise pre-processing such as drying to be done close to where kelp is harvested), many of which are within the territories of Indigenous people and governments. Partnering with Indigenous communities will be essential to ensure the growing industry and its supply chains will benefit, rather than impact, Indigenous waters, communities, and rights. This will require a just regulatory environment to ensure development proceeds in an equitable, rights-driven manner ^[46,57,58]^. There is also potential for conflict between seaweed aquaculture and other marine industries (e.g., tourism, shipping, and fisheries) if the industry should develop to the extent envisioned by our more ambitious scenarios.

Navigating these conflicts, possibly through marine spatial planning, will be essential ^[54,59]^.

Similarly, the potential positive and negative ecological effects of expansive seaweed aquaculture are myriad. Seaweed aquaculture can improve water quality ^[60,61]^, protect shorelines ^[4,62]^, create refugia from ocean acidification ^[4,63]^, and provide habitat and nutrients to various marine species ^[61,64–66]^, however these benefits will be context and species dependent ^[42]^. On the other hand, the large areas required for effective CDR may lead to competition for nutrients and light, reducing productivity of wild seaweeds, phytoplankton, and benthic communities ^[6,47,67]^. There are also risks of harmful algal blooms, non-native species introductions, endemic and emerging pathogens and diseases, and the potential for genetic interactions with wild seaweed populations ^[42,68,69]^. Additionally, marine sequestration of seaweed biomass may negatively impact nearby sediments ^[42,70]^, as well as mesopelagic and deep-sea food webs and water chemistry ^[6,31,71]^. Further research into social, economic and ecological implications such as these will be critical for guiding the development of seaweed aquaculture in Canada, and elsewhere around the world.

Our modelling shows a wide range of plausible results within scenarios, as well as significant differences between them. The first reflects uncertainty in the model parameters, while the latter reflects the uncertain future of seaweed aquaculture development in BC. Exploration of uncertainty using sensitivity analyses identified kelp production rates as a key source of uncertainty. The estimated production rates provided by regional seaweed producers were variable, as well as substantially lower than some global estimates, which resulted in large uncertainties for these model parameters (i.e., a mean of 0.78 kg ww / m^2^ for *Saccharina latissima* vs. an average literature value of 8.3 kg ww / m^2^, see Supplementary Methods for further information). Optimising production rates and accounting for sources of variability in rates of growth and erosion, such as environmental factors, nutrient limitations, and time of crop harvest ^[30,72]^, none of which were considered here, will be important for reducing the uncertainty in the climate benefits of seaweed aquaculture. Production rates in line with average values from the literature were used in two of our scenarios (*Expanded - Optimized* and *Techno Industrial*) to reflect the potential for increased production as the industry in BC develops.

Although less significant than production rates, model estimates of net climate benefit are also sensitive to numerous other sources of uncertainty (e.g., the emissions offset by replacing conventional food products with a seaweed-based alternative, the emissions from production of material used in aquaculture operations, the export of seaweed carbon from aquaculture, and the flux of CO_2_ from the air to surface waters) ^[20]^. Improving our understanding of these various processes, particularly as they apply to local settings, will be critical for accurately quantifying the climate benefits of seaweed aquaculture.

Climate change caused by increasing atmospheric CO_2_ concentrations is a global challenge, however CDR strategies such as kelp aquaculture will be entirely place-based. This can make global assessments of limited use to those on the ground. By this we mean that the success of CDR strategies will depend very much on local environmental and ecological suitability, as well as local production costs and supply chains, community buy-in, and good governance ^[58]^. We therefore advise caution in relying on global models and parameters ^[5,22,73,74]^ to assess local feasibility, as these are unlikely to provide the accuracy necessary to answer essential ecological, economic, and certification questions. To advance assessments of feasibility, we need to transfer such models and their parameterization to local social-ecological contexts.

By integrating seaweed aquaculture industry emissions, in-water sequestration, and emissions avoided through the production of replacement products, this work provides novel insight into the potential role of seaweed aquaculture as a climate change mitigation strategy. Despite the significant uncertainties, our results indicate that kelp aquaculture can contribute to atmospheric greenhouse gas reduction, with values for BC on par with other natural climate solutions examined across Canada, thus helping BC and Canada achieve their climate goals. Realizing this potential will require replacing carbon-intensive products with lower emission, seaweed-based products. Marine sequestration pathways are important to consider, but appear unlikely to have substantial climate benefit and entail potentially significant environmental consequences. Further research to refine our understanding of the pathways modelled here will help advance our understanding of the potential climate mitigation benefits of seaweed aquaculture.

## Methods

### Experimental Design

The mathematical model developed here aims to describe the carbon sequestration potential as well as the associated emissions for seaweed aquaculture and various possible fates for harvested seaweed biomass. The structure of the model was informed by reviewing several published seaweed aquaculture models ^[6,31,37,74,75]^, and consists of 12 equations with a total of 81 parameters. We divided the model into sub-models for seaweed production, carbon sequestration and product fates, and emissions. The production sub-model first estimates the biomass of seaweed that could be produced in the defined study area. We then track the fate of this biomass along the pathways represented in the carbon sequestration and products fate sub-model, and estimate the emissions produced along each pathway with the emissions sub-model. We used the R statical package ^[76]^ to develop our model. Here, we outline the main equations and briefly describe how each was parameterized. Additional details are provided in the Supplementary Methods.

#### Seaweed Production

We calculated the species-specific biomass of kelp produced across the study area as:

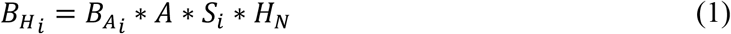

where 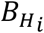 is the total wet weight (ww) of biomass harvested annually for species *i* (kg ww/year); 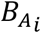 is the area-based kelp harvest for species *i* (kg ww/m^2^); *A* is the total area used for kelp production (m^2^); *S*_*i*_ is the proportion (unitless) of the area used to produce species *i*; and *H*_*N*_ is the number of harvests per year. We included the three kelp species most commonly farmed in BC: *Saccharina latissima* (sugar kelp), *Alaria marginata* (ribbon/winged kelp), and *Nereocystis luetkeana* (bull kelp). The model does not explicitly account for nutrient availability or other factors that may influence production rates. Instead, we used a range of seaweed production estimates obtained from local seaweed producers and the literature to capture the variability in production rates across different environments.

Some portion of kelp biomass produced is lost as detritus prior to harvest in the form of POC and DOC ^[22,23]^. While likely small compared to harvested biomass, this POC and DOC can contribute to sequestration. We therefore related POC and DOC portions to *B*_*H*_ by first back-calculating the net primary productivity (NPP) as:

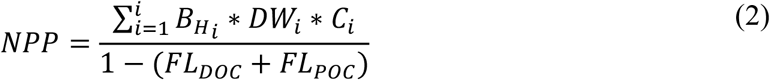

where *NPP* is in (kg C/year); 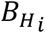 is the total harvested biomass for species *i* (kg ww/year); *DW*_*i*_ is the wet- to dry-weight conversion for species *i* (kg dw / kg ww); *C*_*i*_ is the carbon content of species *i* (kg C / kg dw); *FL*_*DOC*_ is the estimated fraction of kelp carbon lost as DOC (unitless); and *FL*_*POC*_ is the estimated fraction of kelp carbon lost as POC (unitless).

With NPP estimated, we then calculated POC and DOC (in kg C/year) as:

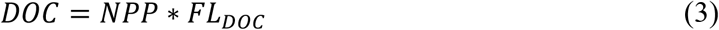

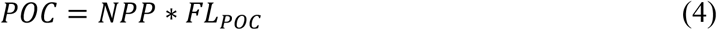

where *FL*_*DOC*_ and *FL*_*POC*_ are as above. We obtained estimates of the carbon content of seaweed from Duarte ^[77]^, and the fractions of DOC and POC lost from Krause-Jensen and Duarte ^[22]^.

#### Sequestration and Product Replacement

We estimated the potential for carbon sequestration (in kg CO_2_/year) as the sum of the sequestration and emission avoidance pathways:

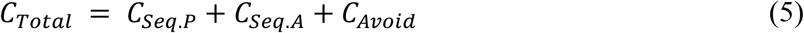

Total carbon sequestration (*C*_*Total*_) is the sum of passively sequestered carbon (*C*_*Seq*.*P*_) via POC and DOC, actively sequestered carbon (*C*_*Seq*.*A*_) via purposefully leaving or releasing harvested kelp into the marine environment, and carbon emissions avoided by replacing other products (*C*_*Avoid*_). Carbon sequestration values are expressed as kg CO_2_/year. Calculation of the sequestration related values include a correction to account for the biological (e.g., respiration) and oceanographic (e.g., upwelling) processes which replace CO_2_ in surface waters and means the carbon sequestered in seaweed tissue does not have a one to one relationship to atmospheric drawdown of CO_2_ ^[47,74]^. Details on each of the pathways in Equation 5 and the associated parameters can be found in the Supplementary Methods.

#### Emissions

We estimated total carbon emissions (in kg CO_2_e/year) from the production and use of kelp as:

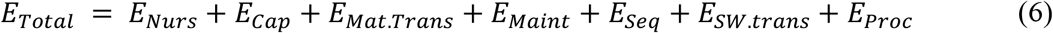

Total carbon emitted (*E*_*Total*_) is the sum of emissions from nursery operations (*E*_*Nurs*_), the production of capital equipment (*E*_*Cap*_), material transport (*E*_*Mat*.*Trans*_), farm maintenance (*E*_*Maint*_), active sinking of kelp (*E*_*Seq*_), the transport of harvested kelp to port (*E*_*SW*.*trans*_), and the processing of kelp into final products (*E*_*Proc*_). All emissions are expressed as kg CO_2_e/year. This sub-model captures many of the primary sources of emissions, however other potentially important emissions such as those from waste management, product storage, or product transport could not be included due to insufficient information. We provide the details on the representation of the pathways underlying the emissions in Equation 6 and the associated parameters in the Supplementary Methods.

### Parameter Values

We obtained parameter values from the literature and through discussions with seaweed producers in BC and Alaska. In this region, the seaweed cultivars are dominated by fast-growing brown kelps (order Laminariales, including *Saccharina latissima, Alaria marginata, Nereocystis luetkeana, and Macrocystis spp*.). Kelp aquaculture in this region typically involves culturing kelp ‘seed’ (gametophytes and juvenile sporophytes) in a controlled nursery, which is then applied to floating longlines for a single cultivation and harvest period each year. Harvested kelp biomass is transported to primary processing (e.g., freezing, drying) facilities by boat, after which the biomass may travel onwards for additional processing.

Wherever possible, we gave preference to parameter values from local seaweed producers or literature values derived in the eastern North Pacific. For each parameter we defined a quantitative distribution when data were sufficient to provide a standard deviation or a minimum and maximum value. When data were insufficient, uncertainty was assessed qualitatively to define a distribution (e.g., ± 50%). We used normal (often truncated at zero), uniform, or triangular parameter distributions depending on the available information. Further discussion of parameter values and how they were derived is provided in the Supplementary Material.

For the model parameters that vary spatially (e.g., transport emissions between port and the farm site) we used a zonal approach. We calculated these parameters using area-weighted averages to account for the envisioned extent of kelp farms under each scenario (see building kelp farming scenarios below and Supplementary Material).

### Engagement with Regional Kelp Producers

We engaged with kelp producers from California to Alaska to parameterise the model and develop appropriate scenarios. We conducted unstructured interviews and distributed a questionnaire (see Supplementary Table S6), with a focus on kelp production rates, and product and emission pathways. We received five responses with varying levels of detail. Some respondents declined to answer specific questions for proprietary reasons, while several provided detailed responses regarding production and maintenance. Less information was provided on emissions as data were often not available or hard to access. One individual declined to participate because the level of detail requested was too specific.

### Building Kelp Farming Scenarios

Our first two scenarios characterize the state of affairs reported by current producers in BC, expanded to 507 km^2^ of suitable area in shallow waters, and assuming farms remain close to coastal communities. The first scenario (*Local – No Harvest*) assumes all kelp is left in the water, akin to natural kelp beds. We included this scenario as a point of comparison for the remaining scenarios. The second scenario (*Local - Products)* is identical to the first in terms of spatial extent and production, but assumes kelp is harvested and used for various purposes as reported by local producers (Table 1).

Our third (*Expanded*) and fourth (*Expanded – Optimized*) scenarios represent expansion of the industry to 1,210 km^2^ of shallow and deep waters in close proximity to communities. Kelp fates for both scenarios resemble those used in the *Local* scenarios but are further diversified to represent emerging new markets for seaweed biomass (e.g., perhaps driven by the saturation of existing markets; Table 1). The *Expanded* scenario uses the kelp production rates reported by producers (as with the first two scenarios), while the *Expanded – Optimized* scenario uses production rates from the literature which are substantially higher than those currently reported by producers in BC and Alaska. The literature values include production rates from field studies and modelling in various temperate locations (see Supplementary Material), and reflect the potential for optimization as the industry develops.

The fifth and final scenario (*Techno Industrial*), represents a maximal approach, assuming kelp farming covers 5,681 km^2^ extending to deep and shallow areas across the coast, regardless of proximity to communities. In this scenario, half of all kelp produced is transported and sunk in deep water with the remainder for replacement products (Table 1). As with the *Optimistic Future* scenario, production rates are assumed to be optimized, and are based on values reported in the literature.

### Spatial Extent of Kelp Aquaculture

To estimate the area available for kelp aquaculture in BC under each scenario we used suitability restrictions based on depth, substrate, proximity to communities, and existing human uses. All spatial calculations were performed using ArcGIS 10.3 ^[78]^.

Current kelp farms in BC and Alaska have a small footprint (up to 8 hectares or 20 acres each) and are largely focused on value-added production. They are generally limited to shallower depths and soft or mixed substrates to facilitate anchoring, and also tend to be located close to coastal communities to facilitate logistics (*Local – No Harvest* and *Local – Products* scenarios). We assumed that with additional investment, farms could operate in deeper waters (though with higher emissions), greatly expanding the potential footprint on the coast both in close proximity to communities (*Expanded* and *Expanded - Optimized* scenarios) and in more remote areas (*Techno Industrial* scenario).

We identified suitable depths using a 100 m bathymetry ^[79]^. We excluded areas less than 15 m to ensure sufficient farm depth, and greater than 200 m (the approximate depth of the shelf break). Shallow waters were defined as between 15 and 75m depth, while deeper waters were 75-200 m. We identified suitable substrate based on Gregr et al. ^[80]^ and defined all soft-bottom areas as suitable. To represent proximity to existing coastal communities, we buffered populated locations ^[81]^ to 25 km to define marine zones in close proximity to communities. Depth, substrate, and proximity restrictions were applied based on feedback from local producers.

We estimated the size of incompatible human use areas (i.e., those dominated by transportation, commercial fishing, recreation, and protected areas) based on footprints of cumulative impacts in the coastal environment ^[82]^. We defined areas incompatible with kelp farming as those with existing human uses above a minimum cumulative effect score. For the coastal footprint, we selected a minimum threshold (2.5) as this provided some exclusion around larger coastal communities in BC.

In addition to these spatial restrictions, we further limited kelp aquaculture to 10% of the total available area to account for additional spatial restrictions not reflected in our calculations (e.g., wave exposure, nutrient limitation, additional competing uses) and the significant logistical challenges faced by many areas of BC’s remote coast. Other modelling efforts have made similar but less conservative assumptions (e.g., Spillias et al. ^[40]^ assume 50% suitability). Our approach is intentionally conservative, reflecting the important environmental conditions and local contexts not accounted for in our calculations.

### Sensitivity Analysis

We used a sensitivity analysis to assess the relative contribution of different model parameters to the uncertainty in our model estimates. We assessed model sensitivity at the resolution of each sub-model (Production, Sequestration and Products, Emissions) and for the individual parameters within each sub-model. We assessed sensitivity using Monte Carlo simulations where the parameter(s) of interest were sampled with uncertainty while holding all other parameters at their central estimate. We used 10,000 runs to describe sub-model sensitivities and 1000 runs for individual parameter sensitivities.

We were primarily interested in the effect of parameter uncertainty on the predicted net climate benefit (Tg CO_2_e / year). We converted this to a relative value by dividing the estimate from each sensitivity analysis by the predicted net climate benefit with no uncertainty (i.e., with all parameters held at their central estimate). This allowed comparisons between sensitivity analyses as well as to the full model, providing insight into the uncertainty in each sub-model as well as in specific parameters. Sensitivity analysis results are scenario-specific, varying based on the parameters used and the selected kelp fates.

## Acknowledgements

We are grateful to Conelia Rindt, Heidi Alleway, and Tiffany Waters who shared their expansive knowledge of seaweed aquaculture with us, provided invaluable feedback, and made this project possible. We also acknowledge Louis Druehl, Lee-Ann Ennis, Majid Hajibeijy, and Jennifer Clark for providing information on their kelp farming operations. We thank The Jordan and Andrea Lott Foundation, Nature United, and The Nature Conservancy for funding this work.

## Author Contributions

Conceptualization: JD, EG

Methodology: CB, EG, JD, JB, TS, MHL

Investigation: : CB, EG

Visualization: CB, JD, EG

Supervision & project administration: JD, JB, EG

Writing—original draft: EG, CB

Writing—review & editing: CB, JD, EG, JB, TS, MHL

## Data Availability Statement

All data are available in the main text or the supplementary materials.

