## Supplementary materials for "Climate benefits of seaweed farming: estimating regional carbon emission and sequestration pathways"

Cameron Bullen *et al.*

**This PDF file includes:**

Supplementary Methods
Supplementary Figures. S1 to S4
Supplementary Tables S1 to S6
References

**Supplementary Methods**

Here we provide additional details on the equations used to calculate production, sequestration, and emissions components of the model. The equations use a number of parameters to scale production, emission, and sequestration values. We also provide detailed descriptions of parameter values and how they were obtained.

Seaweed Production

We converted the data obtained on seaweed production to a generalizable area-based estimate of production:

Equation S1: $$B_A = \frac{B}{H_N} * A_f$$

Where $B_A$ is the area-based seaweed harvest (kg ww/m$^2$); $B$ is the total harvested biomass (reported by growers or found in the literature; kg ww/year); $H_N$ is the number of harvests per year, and $A_f$ is the total area farmed (m$^2$).

We calculated area-based biomass harvest estimates for each species of seaweed farmed in BC. Where multiple estimates were available for a given species, we used the mean and range of the estimates to create a distribution for the parameter (Table S4).

Sequestration and product replacement pathways

Passive carbon sequestration reflects the carbon sequestered during the growing period via the creation of particulate and dissolved organic carbon (POC and DOC), some fraction of which is sequestered. We calculated this as:

Equation S2: $$C_{Seq.P} = [(POC * F_{POC.Seq} * Seq.Mult) + (DOC * F_{DOC.Seq} * Seq.Mult)] * K_{C.CO2} * K_c$$

Where $POC$ and $DOC$ are the amount of POC and DOC produced, calculated by the production sub-model (kg C/year); $F_{POC.Seq}$ is the fraction of the POC sequestered (unitless); $Seq.Mult$ is a multiplier used to model sequestration differences associated with farm depth; $F_{DOC.Seq}$ is the fraction of the DOC sequestered (unitless); $K_{C.CO2}$ is the carbon to $CO_2$ conversion (kg C/kg $CO_2$); and $K_c$ is the conversion factor that relates carbon sequestered to atmospheric drawdown of $CO_2$.

Active carbon sequestration describes the sequestration from purposeful release of harvested seaweed into the marine environment. This can consist of either release near the farm or transport to specific locations and active sinking of harvested biomass where the released kelp is more likely to reach long-term storage (i.e., depths > 2000 m). We calculated carbon sequestration from these two approaches as:

Equation S3: $$C_{Seq.A} = B_{H_i} * DW_i * C_i * F_{Active.Seq} * K_{C.CO2} * K_c$$

Where $B_{H_i}$ is the total harvested biomass for species $i$ (kg /year); $DW_i$ is the wet-weight to dry-weight conversion of species $i$ (kg dw / kg ww); $C_i$ is the carbon content of species $i$ (kg C / kg dw); $F_{Active.Seq}$ is the fraction of actively released seaweed that is sequestered (unitless) $K_{C.CO2}$ is the carbon to $CO_2$ conversion (kg C/kg $CO_2$); and $K_c$ is the conversion factor that relates carbon sequestered to atmospheric drawdown of $CO_2$. We varied the value of $F_{Active.Seq}$ depending on whether the seaweed is released at the site of production or transported to a more favourable location.

The use of seaweed products as alternatives to traditional products such as food, animal feed, or fuels results in the avoidance of carbon emissions associated with the production of these traditional products. We calculated these avoided emissions as

Equation S4: $$C_{Avoid} = B_{H_i} * DW_i * F_{Rep}$$

Where $B_{H_i}$ is the total harvested biomass for species $i$ (kg /year); $DW_i$ is the wet-weight to dry-weight conversion of species $i$ (kg dw / kg ww); and $F_{Rep}$ is the factor relating the carbon emissions avoided for each kg dw of seaweed, and varies by each product class (kg $CO_2$/kg dw).

Emission pathways

This pathway captures the emissions from the operation of a nursery to produce seeded line for seaweed production. We assume that the majority of these emissions are due to energy use, so calculate nursery emissions as:

Equation S5: $$E_{Nurs} = NRG_{Nurs} * A * E_{Nrg}$$

Where $NRG_{Nurs}$ is the annual energy required to run the nursery (GWh / km$^2$ / year); $A$ is the total area used for seaweed production (km$^2$); and $E_{Nrg}$ is the emissions associated with energy production (kg $CO_2$ / GWh).

We calculated capital emissions from the production of the materials needed as part of the capital investment in a seaweed farming operation as:

Equation S6: $$E_{Cap} = \frac{E_{Mat} * A}{Mat.Life}$$

Where $E_{Mat}$ is the average emissions associated with the material required by a square kilometre of seaweed aquaculture (kg $CO_2$ / km$^2$); $A$ is the total area used for seaweed production (km$^2$); and $Mat.Life$ is a multiplier reflecting spatial differences in the expected amount and lifespan of equipment required by seaweed aquaculture at different depths.

We calculated emissions from the transport of material (e.g., ropes, anchors, buoys, etc.) to and from the site of seaweed aquaculture from the nearest port as:

Equation S7: $$E_{Mat.Trans} = M_{Mat} * A * E_{Barge} * 2 * D_{Port}$$

Where $M_{Mat}$ is the mass of equipment required per harvest for each $km^2$ of seaweed aquaculture ($kg / km^2$); $A$ is the total area used for seaweed production ($km^2$); $E_{Barge}$ is the emissions from transporting one kg of equipment one kilometre by barge (kg $CO_2$ / kg / km); and $D_{Port}$ is the distance from the port to the site of farming (km).

We calculated emissions due to maintenance of the kelp farm as:

Equation S8: $$E_{Maint} = ((2 * D_{Port} * E_{Ves} * (\frac{N_{Maint}}{A_{Maint}/A}) * H_N) + (A * D_{Maint} * E_{Ves}))$$

Where $D_{Port}$ is the distance from the port to the site of farming (km); $E_{Ves}$ is the emissions from a maintenance vessel travelling one kilometre (kg $CO_2$ / km); $N_{Maint}$ is the number of maintenance trips required per harvest; $A_{Maint}$ is the area maintained per trip ($km^2$); $A$ is the total farmed area ($km^2$); $H_N$ is the number of harvests per year; and $D_{Maint}$ is the distance travelled per $km^2$ maintained ($km / km^2$).

Active sequestration of seaweed will entail emissions due to transporting seaweed to the optimal sinking location and the sinking process itself. We calculated this as:

Equation S9: $$E_{Seq} = (B_H * E_{Barge} * D_{Sink}) + (B_H * E_{Sink})$$

Where $B_H$ is the amount of harvested seaweed biomass (kg/year); $E_{Barge}$ is the emissions from transporting one kg of equipment one kilometre by barge (kg $CO_2$ / kg / km); $D_{Sink}$ is the distance from the farm to the location of sinking (km); and $E_{Sink}$ is the emissions generated during the process of sinking the seaweed (kg $CO_2$ / kg).

To produce kelp products the harvested seaweed must be transported to the nearest port or processing location. We calculated these transport emissions as:

Equation S10: $$E_{SW.trans} = B_H * E_{Barge} * D_{Port}$$

Where $B_H$ is the amount of harvested seaweed biomass (kg/year); $E_{Barge}$ is the emissions from transporting one kg of equipment one kilometre by barge (kg $CO_2$ / kg / km); and $D_{Port}$ is the distance from the port to the site of farming (km).

Once at port the processing of seaweed into products will also entail greenhouse gas emissions. In the absence of details on emissions associated with the different potential products, we calculated these emissions simply as:

Equation S11: $$E_{Proc} = B_H * E_{Conv}$$

Where $B_H$ is the amount of harvested seaweed biomass (kg/year); and $E_{Conv}$ is the emissions released from converting seaweed into a final product.

Parameter values

Here, we provide a detailed description of the values used for all parameters in the model, with a summary of these values provided in Table S3.

**Species-level productivity data**
*Regional values*
We collected kelp production rates for species used in aquaculture operations in BC, Washington, and Alaska from local producers  and the published literature. Overall, we heard that the majority of aquaculture operations are growing Saccharina, with some limited production of Alaria and Nereocystis. Macrocystis was originally included in the model but was not reported to be grown commercially in BC. From the values obtained, we calculated a mean area-based production rate (kg ww / m2 / harvest) and a standard deviation for each species (Table S4). Monte Carlo runs were drawn from truncated normal distributions (a normal distribution constrained to be > 0).

*Literature values*
We also compiled production values reported in the literature and used in other seaweed aquaculture models to represent production rates that have been achieved elsewhere and could, in theory, be achieved in BC (Table S5). We compiled values for a variety of temperate brown seaweeds, and used the mean and standard deviation of these compiled values in the *Expanded - Optimized* and *Techno Industrial* scenarios.

**Total Farmable Area (A)**
We calculated the total area suitable for seaweed aquaculture on the BC Coast by scenario as described in the main text.

**Area-Based Biomass ($B_A$)**
These values are species specific, and were calculated from either reported harvest rates for the Pacific Coast of North America (*Local - No Harvest, Local - Products*, and *Expanded* scenarios) or from values reported in the literature (*Expanded - Optimized* and *Techno Industrial*). See Table S3 and S4 for details and specific values.

**Species Composition (S)**
Three species of seaweed are currently farmed in BC: bull kelp (*Nereocystis leutkeana*), ribbon kelp (*Alaria marginata*), and sugar kelp (*Saccharina latissima*). For the purpose of this model, we presumed seaweed production followed the approximate composition of current kelp aquaculture in BC which is dominated by Saccharina. We therefore assigned 80% of the total farmable area to grow Saccharina, with 10% allocated to each of Alaria and Nereocystis. Macrocystis was initially considered for the model but was dropped as no aquaculture operations reported farming this species commercially in BC.

**Number of Harvests ($H_N$)**
We included the number of harvests in the model to reflect the possibility for multiple seaweed harvests per year. We heard from local seaweed producers that seaweed is currently harvested

once a year in the spring, but investigations into the potential for a second harvest later in the year are ongoing. For all model runs the number of harvests was set as 1.

**Dry-Weight Conversion (DW)**
We obtained wet-weight to dry-weight conversions for each species from the literature. Where different estimates were available for blades, stipes, or mixed samples, we selected mixed estimates as this assumes both blades and stipes will be harvested. We assumed these conversion values to be constant, with no uncertainty.

*Nereocystis*: 0.129 kg dw / kg ww [1]
*Alaria*: 0.165 kg dw kg dw / kg ww [1]
*Saccharina*: 0.113 kg dw / kg ww [2]

**Carbon Content (C)**
We obtained carbon content of seaweed from the literature. We used the estimated mean of 0.248 kg C / kg dw for reported global macroalgae [3], and the reported standard deviation of 0.063 kg C / kg dw to capture the uncertainty in this estimate.

One survey respondent reported carbon contents of 0.325 and 0.28 kg C / kg dw for Alaria and Saccharina, respectively, for low wave-energy aquaculture sites. These estimates are similar to, but greater than the mean estimate from Duarte [3]. As they fall within the range of values used to parameterise the model, we retained the values from Duarte [3].

**Seaweed Lost as DOC ($FL_{DOC}$)**
We calculated the fraction of seaweed lost as DOC based on the global estimates provided by Krause-Jense and Duarte [4]. We calculated the central estimate using the reported median values for global production (1.419 PgC /yr) and DOC export (0.330 PgC/yr) yielding an estimate of 23.3% (or 0.233 kg DOC / kg C).

We characterized the uncertainty around this estimate by calculating maximum and minimum values for DOC export based on the reported 25th and 75th percentiles [4]. The minimum and maximum values were calculated to be 13.7% and 34.2%, respectively.

Reed et al. [5] estimated a DOC production rate of ~ 8 mg C / g dw / day for Macrocystis forests in southern California. Roughly assuming a growing season of 100 days and a seaweed carbon content of 0.248 kg C / kg dw, we arrive at an estimate of 0.32 kg DOC / kg C. This value is higher than the median estimate from Krause-Jense and Duarte [4], but falls within the range of expected values. We therefore retained the estimates from Krause-Jense and Duarte [4] for use in the model.

**Seaweed Lost as POC ($FL_{POC}$)**
We calculated the fraction of seaweed lost as POC in a similar fashion based on the global estimates provided by Krause-Jense and Duarte [4]. We calculated the central estimate to be 16.7%. We calculated minimum and maximum values based on the 25th and 75th percentiles to be 0% and 55% (note: the 25th percentile reported by Krause-Jense and Duarte [4] is negative, but we set it to zero here as the lowest possible value).

**DOC Sequestered ($F_{DOC.Seq}$)**
We calculated the fraction of DOC sequestered based on estimates from Filbee-Dexter et al. [6]. Filbee-Dexter et al. [6] used decomposition rates and a hydrodynamic model to generate a coarse estimate for the amount of detrital production (DOC and POC) sequestered at various sites around the globe, of which three were in BC. At these three sites, they estimated that 10.6%, 14.0%, and 40.2% of detrital production, respectively, is exported to deep water (> 200 m) where it can be sequestered. We used these three estimates to define a triangular distribution with a minimum of 10.6%, a central estimate of 14.0%, and a maximum of 40.2%.

**POC Sequestered ($F_{POC.Seq}$)**
We used the same estimates and uncertainty distribution as above to estimate the fraction of POC sequestered.

**Passive Sequestration Multiplier (*Seq.Mult*)**
In addition to the seaweed detritus generated and sequestered as POC and DOC in shallow waters (i.e., < 50 m, described above), we also considered the effects of farm location, as the efficiency of carbon sequestration is known to vary by depth [7,8]. We therefore defined a multiplier to reflect that seaweed lost from farms in deeper water would be more likely to be sequestered at depth than stay in shallow water or wash up on shore. We applied this multiplier only to the amount of POC and DOC sequestered, not to the amount of seaweed lost. Although it is conceivable that increasing depth may be associated with increased wave exposure and therefore increase detrital production.

In the 15-75 m depth class, we used a value of 1 (i.e., no multiplier), while in the 75-200 m depth class we sampled the multiplier from a uniform distribution centered on 1.2 (20% increase), with minimum and maximum values set at ±50% (1.1 and 1.3, respectively).

**Seaweed Sequestered when Sunk ($F_{Active.Seq}$)**
The fraction of sunk seaweed sequestered for meaningful time scales (i.e., > 100 years) will depend on the depth to which the seaweed was sunk [8]. Carbon sequestered in marine waters also suffers from leakage, with local circulation and depth playing a significant role in how long carbon will be sequestered [8,9]. For our model we assumed sinking occurs either in shallow water near the site of production (< 200 m) or deep water (> 500 m) and other sites with longer residency times (e.g., marine canyons, fjords).

For deep seaweed disposal (> 500 m), we estimated the fraction of seaweed sequestered based on sequestration curves developed by Siegel et al. [8]. Based on the pixels representing the BC coast on Siegel et al.'s global map of results, we used the 100+ year sequestration proportions for 500 m, 1000 m, and 1500 m depth as the minimum, central, and maximum estimates, respectively. This resulted in a central estimate of 47%, a minimum of 10%, and a maximum of 97%.

For shallow seaweed disposal near farms, we estimated the fraction of seaweed sequestered using values Filbee-Dexter et al. [6] for DOC and POC sequestration as above: once again, we used a triangular distribution based on three estimates for sites in BC, with a minimum of 10.6%, a central estimate of 14.0%, and a maximum of 40.2% of POC sequestered. We also used the sequestration multiplier (Seq.Mult) to account for the role of farm depth in sequestration rates in the two different depth classes considered.

**Carbon to $CO_2$ Conversion ($K_{C.CO2}$)**
We converted units of carbon to units of $CO_2$ using a conversion factor based on the molar mass of C (12 kg/mol) and $CO_2$ (44 kg/mol). This conversion factor is 3.67 kg $CO_2$/kg C.

**Atmospheric Removal Fraction ($K_C$)**
This parameter represents the relationship between the carbon sequestered via seaweed biomass and the drawdown of atmospheric carbon. This parameter is important and complex, as this relationship depends on many factors. It is very sensitive to both biological (e.g., carbon recycling within seaweed beds, competition between seaweed and phytoplankton) and oceanographic factors (e.g., non-instantaneous equilibrium between air and water gas exchange, ocean circulation), leading to a dynamic rate of reduction in atmospheric carbon [10,11]. Berger et al. [11] estimate that macroalgae carbon sequestration draws down 40-80% of its carbon via the ocean from the atmosphere, depending on nutrient levels. Similarly, DeAngelo et al. (2022) estimate the atmospheric removal fraction to be 50%, and for their model use a range of 40-100%.

Based on these estimates we used 50% as our central estimate, with a minimum and maximum value of 40% and 80%.

**Product Replacement Fraction (FRep)**
This parameter captures the carbon emissions avoided for each kg dw of seaweed, and varies by each species and product class. We evaluated three possible end products: food, animal feed, and fuel. We discussed fertilizer and biochar with seaweed producers and found that there is as yet no market for these products in the eastern North Pacific. The replacement fraction generalizes information on the average emissions for a typical product in each category, and estimates the equivalency between the traditional and replacement products (i.e., how many kg of seaweed are required to produce an equivalent product).

*Food*
In their global analysis, DeAngelo et al. [10] estimated GHG emissions avoided for food ranged from 1-6 kg CO2/kg dw. This is based on comparing caloric content of seaweed products to traditional food items and the emissions from the land use and nutrient management of these non-seaweed products (pulses, vegetables, fruits, oil crops, and cereals; carbon estimates for non-seaweed products from Hong et al. [12]). We used these two values as the minimum and maximum, and their average (3.5 kg CO2/kg dw) to describe the triangular distribution used for the Monte Carlo runs. These avoided emissions refer only to those from land use and nutrient management, and not those associated with fertilizer production, or any other processing, storage, or transport activities.

*Animal Feed*
DeAngelo et al. [10] calculated GHG emissions avoided for animal feed as 2-3 kg CO2/kg dw by comparing on a caloric basis, seaweed products to emissions from non-seaweed products (oil crops, and cereals; carbon estimates for non-seaweed products from Hong et al. [12]). We used

these two values as the minimum and maximum, and their average (2.5 kg CO2/kg dw) to describe the triangular distribution used for the Monte Carlo runs. The use of seaweed as a cattle feed additive to reduce methane production was not modeled in this study.

*Fuel*

DeAngelo et al. [10] estimated GHG emissions avoided for fuel (bioethanol) production ranged from 0.7-1 kg CO2/kg dw. We used these two values as the minimum and maximum, and their average (0.85 kg CO2/kg dw) to describe the triangular distribution used for the Monte Carlo runs.

**Nursery Energy Use ($NRG_{Nurs}$)**

Nursery operations include spore preparation, inoculation to lines, and juvenile maturation. In their techno-economic model, Coleman et al. [13] estimated that a nursery capable of providing all the seeded line required by aquaculture in the state of Maine (~ 56,600 m of grow line) would require 92,328 kWh annually. Conservatively assuming a 3 m spacing between lines, this is equivalent to providing seeded line to aquaculture area of 169,800 $m^2$ (0.169 $km^2$). This breaks down to an annual nursery energy use estimate of 0.544 kWh / $m^2$ / year (or 0.544 GWh / $km^2$ / year). An alternate estimate based on seaweed aquaculture in Sweden [14] estimated a requirement of 0.195 kWh / $m^2$ / year based on a 0.5 Ha case study. We used these two values as minimum and maximum values, along with a central estimate at the midpoint (0.37 kWh / $m^2$ / year) to describe the triangular distribution used for the Monte Carlo runs.

**Emissions from Energy Production ($E_{Nrg}$)**

We obtained average emissions ($CO_2$ equivalents) for BC from energy production from the government's electric grid emission intensity factor [15]. The intensity factor indicates the average carbon emissions for every unit of grid energy used. The 2021 intensity factor was 9.7 t $CO_2$ / GWh (or 9700 kg $CO_2$ / GWh) while the 2020 intensity factor was 40.1 t $CO_2$ / GWh (or 40,100 kg $CO_2$ / GWh). This large year-to-year change partially reflects changes in the methodology used to calculate the intensity factor at the end of 2020 [15]. Emission intensity factors for which data were available (except 2011) fell within this range. We therefore used the 2020 and 2021 emission values as maximum and minimum values of the uniform distribution used for the Monte Carlo runs.

**Emissions from Material Production ($E_{Mat}$)**

This parameter captures the upstream emissions from the manufacture of the equipment needed to produce seaweed. In some cases this can account for the majority of on-farm emissions [16,17]. These materials primarily consist of steel chains, ropes, anchors, and buoys. We compiled estimates for this parameter from the following literature.

In their LCA of an European seaweed aquaculture system, van Oirschot et al. [16] estimated that infrastructure for single layer seaweed aquaculture produces 5,280 kg $CO_2$ equivalents per ton of protein produced, based on a material lifespan of 20 years. Using their calculations of 0.77 T protein / Ha / yr, and a yield of 77 T ww / Ha / yr yields an emission rate of 56.5 kg $CO_2$ / T ww harvested.

In five scenarios of Norwegian seaweed farming at different scales, Koesling et al. [18] estimate that aquaculture infrastructure produces ~7000 kg $CO_2$ equivalents per tonne of protein produced. Using their reported conversion factors (0.15 kg DW / kg WW, 0.1 kg protein / kg DW) and protein extraction efficiency (80%), we arrive at an estimate of 84 kg CO2e / T ww.

Finally in their techno-economic model, Coleman et al. [19] estimated annual emissions from infrastructure for a 1000 acre off-shore seaweed aquaculture system to be 92 tCO2e / yr. Alongside their estimated yield of 4489 T ww / yr, this results in as estimated 20.5 kg $CO_2$ / T ww.

To make use of these values in our model so that emissions scale with industry expansion but not (incorrectly) with increased production efficiency, we converted these values to a per-area basis using the estimated mean yield for Saccharina reported by local seaweed producers (780 T ww / $km^2$; see Farm-Level production). This provided us a minimum estimate of 15,990 kg $CO_2$ / $km^2$ [19], a central estimate of 44,070 kg $CO_2$ / $km^2$ [16], and a maximum estimate of 65,520 kg $CO_2$ / $km^2$ [18]. We used these values to describe the triangular distribution for the Monte Carlo runs.

**Annual Mass of Equipment ($M_{Mat}$)**
The annualized mass of equipment required for seaweed aquaculture was compiled from relevant literature. We needed this value to estimate fuel usage for transport to and from aquaculture locations.

In their LCA of an European seaweed aquaculture system, van Oirschot et al. [16] estimated an annualized equipment mass of 0.1135 ton / 100 m of line. Assuming a 3 m spacing between lines yielded a total mass estimate of 378 kg / $km^2$ / yr. In comparison, Aitken et al. [20] estimated that Macrocystis long-line cultivation in Chile required an annualized equipment mass of 1155 kg / $km^2$ / yr. In the absence of further information, we used these values as minimum and maximum values with a central estimate at the midpoint (766.5 kg / $km^2$ / yr) to describe the triangular distribution used for the Monte Carlo runs.

**Material Lifespan (Mat.Life)**
Equipment mass has been annualized (see above) allowing us to simply set material lifespan to be one year. However, depth, wave exposure, and other environmental factors can contribute to variation in material lifespan [10,18]. Increased farm depth also requires additional anchor chain, rope, and potentially other equipment. We approximated this variability by allowing material lifespan to vary by depth class. For shallow farms in the 15-75 m depth range, material lifespan was set at the default of 1 year. For deeper farms in the 75-200 m depth range, material lifespan was estimated to be slightly shorter at 0.8 year. In both cases, minimum and maximum values were set at ±0.1 years (i.e., 0.9-1.1 for 15-75 m; 0.7-0.9 for 75-200 m) to create the uniform distribution used for the Monte Carlo runs.

**Emissions from Barge Transport ($E_{Barge}$)**
DeAngelo et al. [10] estimated emissions from transporting material by barge to be 0.00003 kg $CO_2$ / kg / km. Similarly in their model, Coleman et al. [19] estimate barge and tugboat emissions to be 0.000026 kg $CO_2$ / kg / km. We used these values as the minimum and maximum, along

with their midpoint (0.000028 kg $CO_2$ / kg / km) to describe the triangular distribution used for the Monte Carlo runs.

**Distance to Port ($D_{Port}$)**
We estimated distance to the nearest port separately for farms within 25 km of a coastal community and those further away. This distinction was based on the spatial context of our study area, and defines the area accessible by a vessel travelling at 9 knots for 1.5 hours. For the near farms (within 25 km of a community) we estimated the mean distance to port from a truncated normal distribution (constrained to be > 0) with a mean of 12.5 km and a standard deviation of 5 km. For the distant farms (further than 25 km) we estimated the mean distance to port as 62.5 km with a standard deviation of 15. We used these values to create the truncated normal distributions used for the Monte Carlo runs.

**Emissions from Vessel Travel ($E_{Ves}$)**
DeAngelo et al. [10] estimated emissions from a maintenance vessel to be 2.36 kg $CO_2$ / km. This estimate is based on a typical fishing maintenance vessel (14 m vessel with two 310 hp engines) travelling at an average speed of 9 knots [10].

Coleman et al. [19] used a value of 2.9 L / km for maintenance vessel travel in their technoeconomic model. By combining this with the density of diesel (0.83 kg / L) and diesel emissions (3.2 kg $CO_2$ / kg fuel) we obtained a value of 7.72 kg $CO_2$ / km. We used two values were used as the minimum and maximum, and their midpoint (5.05 kg $CO_2$ / km) to describe the triangular distribution used for the Monte Carlo runs.

**Number of Maintenance Trips ($N_{Maint}$)**
DeAngelo et al. [10] and Aitken et al. [20] used a value of 6 trips / km$^2$ / year and we used this value in our analysis. Without further information on the range of possible values or some indication of uncertainty, we deemed confidence to be low and so used a uniform distribution with minimum and maximum value were set at ± 50% of this estimate (i.e., 3–9 trips).

**Area Maintained per Trip ($A_{Maint}$)**
Following Aitken et al. [20], we assumed a maintenance rate of 0.5 km$^2$ / trip. Without further information we deemed confidence to be low and so used a uniform distribution with minimum and maximum values set at ± 50% of this estimate (0.25 – 0.75 km$^2$ / trip).

**Distance travelled per Area Maintained ($D_{Maint}$)**
Aitken et al. [20] assumed that once at the site of production a maintenance vessel would travel 0.5 km to maintain a 1 Ha farm. Scaling this to farm size by area suggest 50 km travelled to maintain one km$^2$ of farm. We assumed it to be more reasonable to scale according to perimeter (given square farms) yielding an estimate of 5km/km$^2$. We used this as the lower estimate, with 10 km / km$^2$ and 15 km / km$^2$ as the central estimate and maximum, respectively, to describe the triangular distribution used for the Monte Carlo runs.

**Distance to Sinking Location ($D_{Sink}$)**
Distance to an optimal sinking location was estimated separately for the shallow and deep farms (defined above) and was based on the geographic context of our study area. In shallow waters,

we estimated the mean distance to sinking location from a truncated normal distribution with a mean of 100 km and a standard deviation of 35 km. For the deeper farms we used a truncated normal distribution with a mean of 75 km and a standard deviation of 25.

**Emissions from Seaweed Sinking ($E_{Sink}$)**
This parameter captures the emissions from actively sinking seaweed. Active sinking will likely require specific equipment (e.g., weights and baling material) and other energy use leading to emissions. For example, Coleman et al. assume sinking will be achieved using reclaimed concrete as ballast [19].

As this process remains conceptual we found no related estimates of energy use or emissions in the literature. We therefore used a nominal value of 0.00005 kg $CO_2$/kg ww as the midpoint, and allowed our triangular distribution to range from 0 to 0.0001 kg $CO_2$/kg ww. We chose these values after considering the emissions required to convert seaweed into consumer products (see $E_{Conv}$ below). We assumed sinking emissions would be a small fraction (< 1%) of these conversion emissions. We included zero to allow for the possibility that seaweed sinking could be done without producing carbon emissions.

**Emissions from Seaweed Conversion ($E_{Conv}$)**
The emissions for converting seaweed biomass into end products will vary substantially depending on the product, the necessary processing, and how and where the processing is done. Given limited information, we used a single conversion emission parameter for all product types, but considered values from a range of sources and contexts.

Drawing from US department of Energy study [21], DeAngelo et al. [10] estimated the emissions from converting seaweed into products such as food, animal feed, and fuel as 0.0057 kg $CO_2$ / kg dw. Using a conservative wet to dry weight conversion of 0.15 gives an estimate of 0.00086 kg $CO_2$/kg ww.

For comparison, a study on seaweed ethanol production in BC estimated drying emissions would be 0.0224 kg $CO_2$ / kg ww based on an energy requirement of 4 MJ / kg ww and an emissions factor of 5.6 g$CO_2$e / MJ [22]. This estimate does not include the additional emissions from actual ethanol production from the dry seaweed biomass, but we found no suitable values to parameterise this further processing.

Using a Danish case study producing ethanol, protein, and fertilizer, Seghetta et al. [23] estimate drying (6%) and biorefinery operations (25%) jointly account for 31% of all operating emissions. Considering total production of 1273 kg $CO_2$e / Ha and a yield of 10 T ww / Ha, they estimate the emissions from drying and refining to be 0.039 kg $CO_2$e / kg ww.

Finally, using seaweed aquaculture system producing biogas and fertilizer in Sweden, Pechsiri et al. [14] estimated seaweed processing (including packaging, transport, cutting, anaerobic digestion, and biogas upgrading) to emit 0.93 ton $CO_2$e for a 0.5 Ha case study. Given an estimated yield of 11.25 T ww results in estimated emissions of 0.0827 kg $CO_2$e / kg ww.

Since the above estimates span almost two orders of magnitude, we selected the lowest estimate as our minimum (0.00086 kg $CO_2e$ / kg ww; DeAngelo et al. 2022), the highest estimate as our maximum (0.0827 kg $CO_2e$ / kg ww; Pechsiri et al. 2016), and the higher of the two intermediate estimates as our central estimate (0.039 kg $CO_2e$ / kg ww; Seghetta et al. 2016) for the triangular distribution used in our Monte Carlo runs.

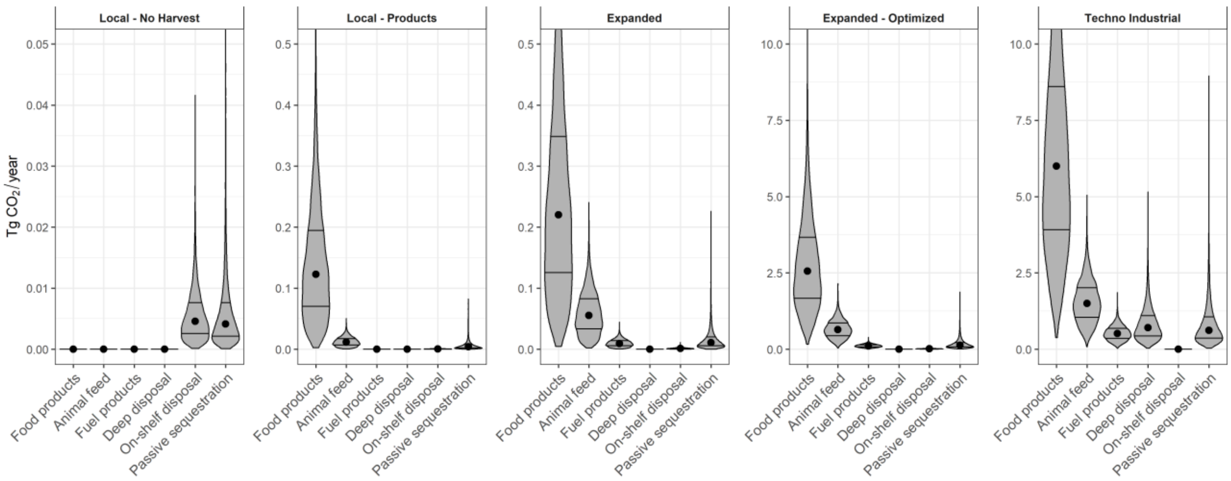


**Fig. S1.**

Atmospheric $CO_2$e reduction potential (Tg $CO_2$ / year), either sequestration or emissions avoided, by pathway for each scenario. The violin plots represent the probability distribution of estimates from 10,000 Monte Carlo runs, while the dot indicates the median and horizontal lines indicate the 25$^{th}$ and 75$^{th}$ percentiles. Note the scale of the y-axis varies between panels.

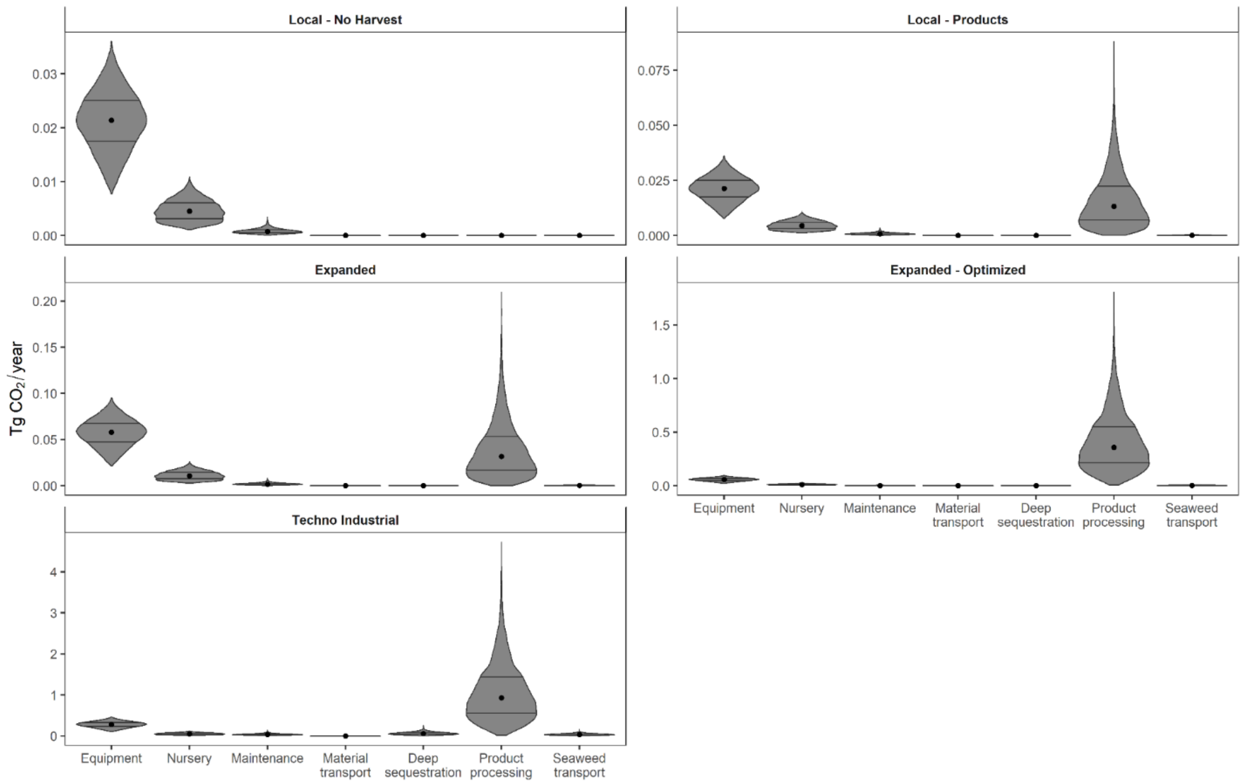


**Fig. S2.**

Emissions (Tg $CO_2$ / year) by sources for each scenario. The violin plots represent the probability distribution of estimates from 10,000 Monte Carlo runs, while the dot indicates the median and horizontal lines indicate the 25$^{th}$ and 75$^{th}$ percentiles. Note the scale of the y-axis varies between panels.

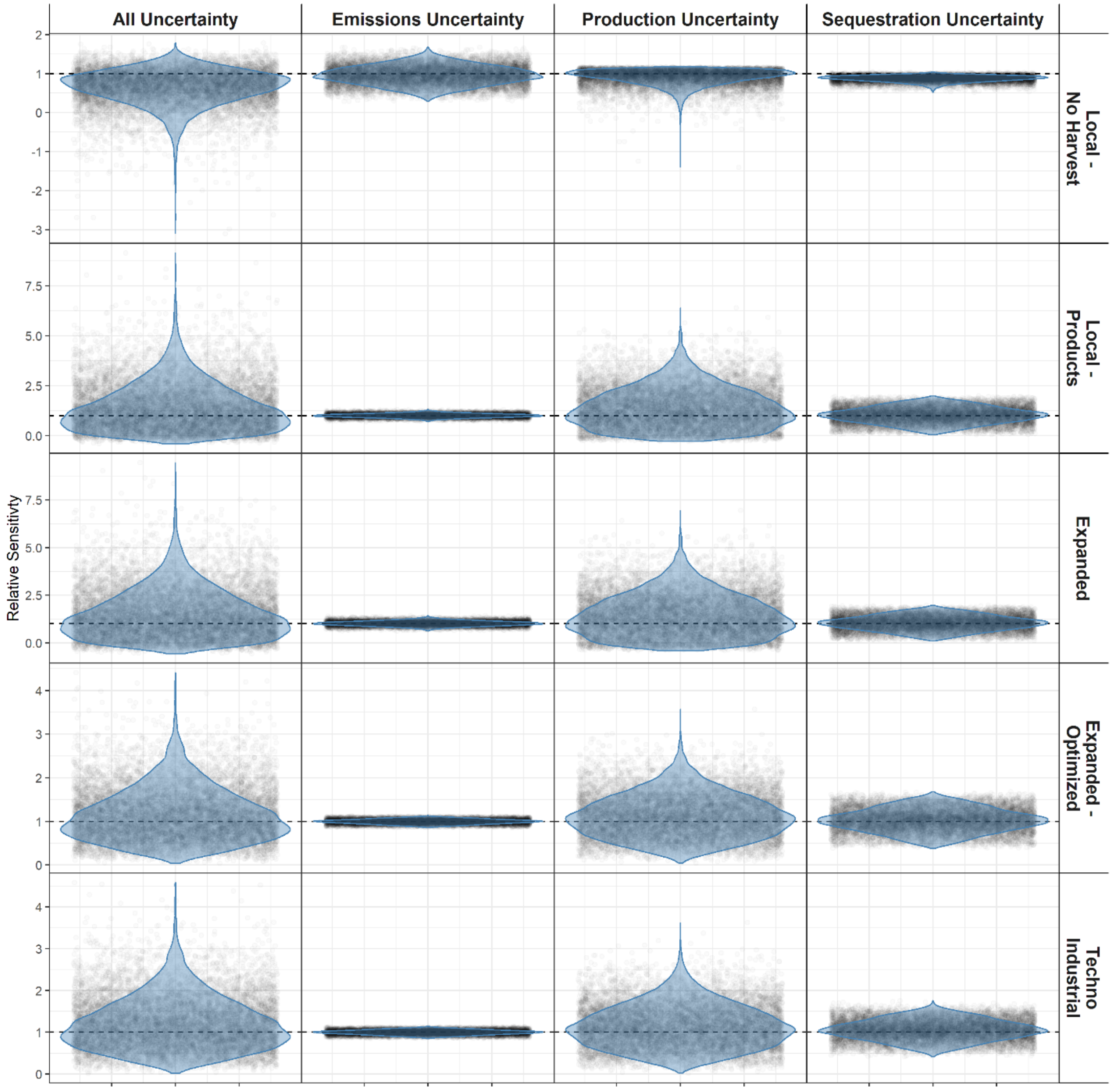


**Fig. S3.**

Relative sensitivity of the estimated net reduction in atmospheric $CO_2$ to cumulative uncertainty in different parameter categories for each scenario. Sensitivity is assessed relative to a model run with no uncertainty, in which model parameters are set at their central estimate. A value of 1 indicates the estimated net reduction in $CO_2$ is the same as a model run without uncertainty, while a value of 0.5 or 2 would represent a halving or doubling of estimated climate benefits, respectively. Violin plots show the distribution of estimates from all Monte Carlo runs, with each individual estimate shown as a shaded point. Note the scale of the y-axis varies between panels.

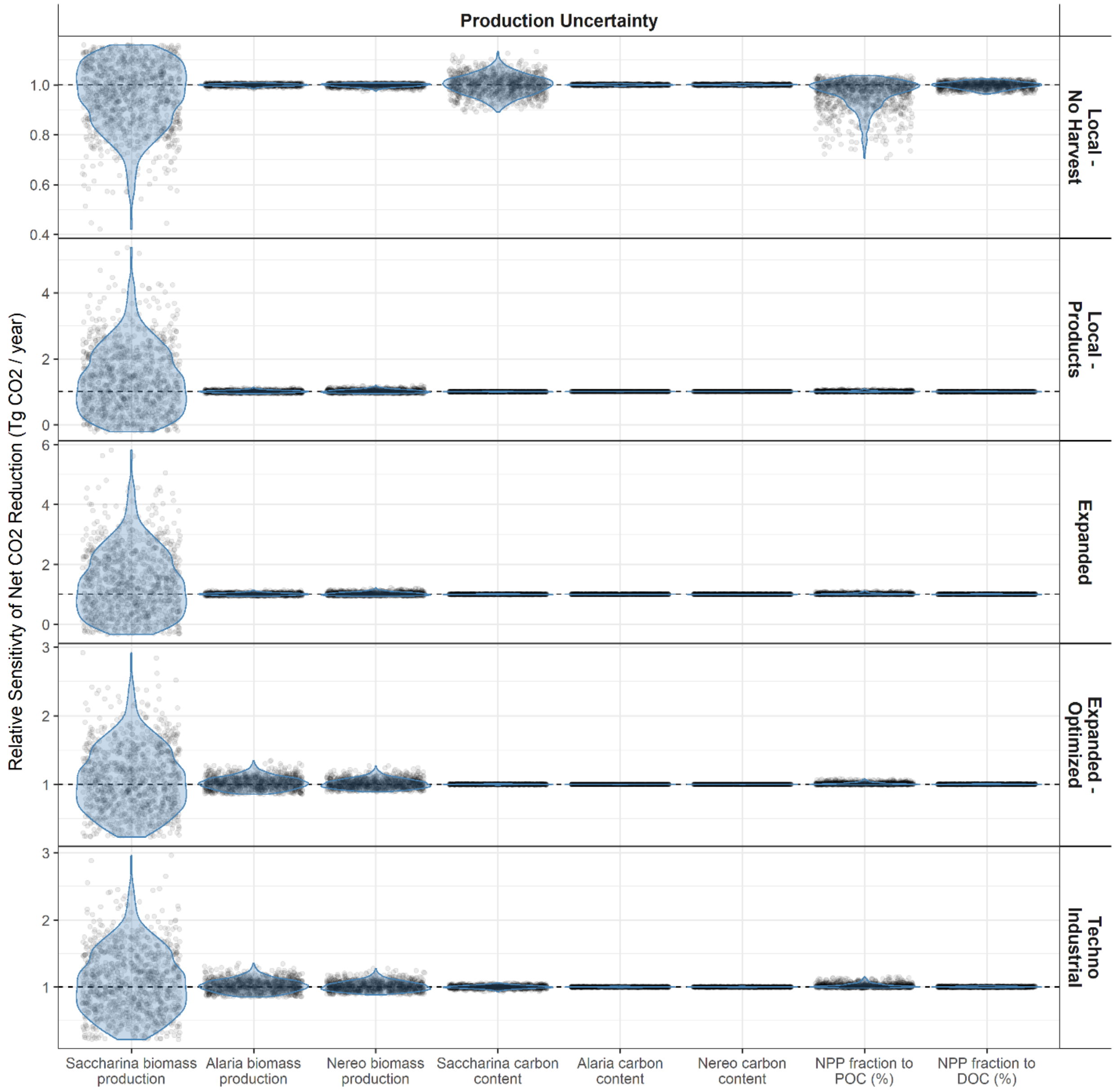


**Fig. S4A.**

Relative sensitivity of the estimated net reduction in atmospheric $CO_2$ to uncertainty in key production parameters for each scenario. Sensitivity is assessed relative to a model run with no uncertainty, in which model parameters are set at their central estimate. A value of 1 indicates the estimated net reduction in $CO_2$ is the same as a model run without uncertainty, while a value of 0.5 or 2 would represent a halving or doubling of estimated climate benefits, respectively. Violin plots show the distribution of estimates from all Monte Carlo runs, with each individual estimate shown as a shaded point. Note the scale of the y-axis varies between panels.

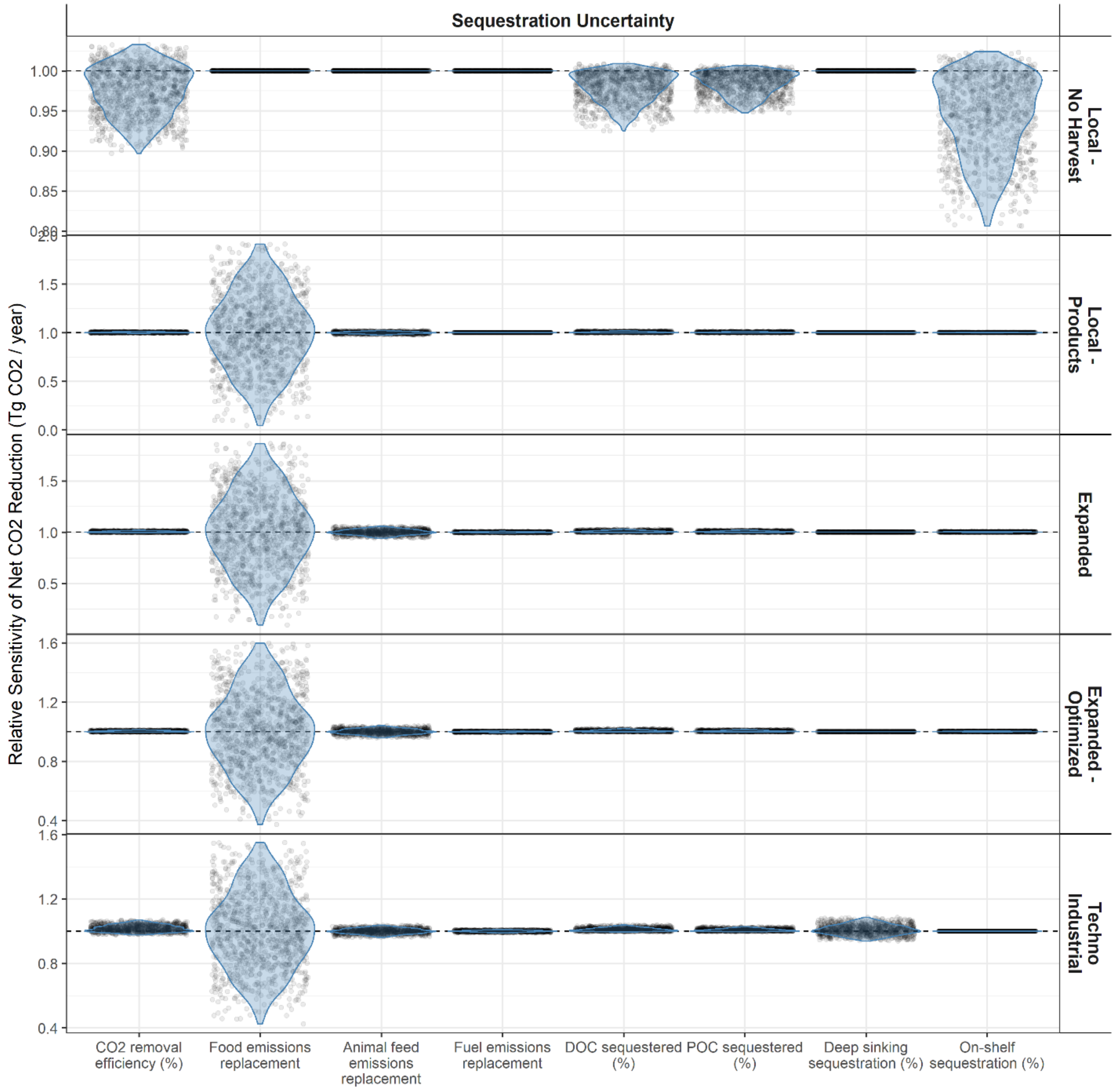


**Fig. S4B.**

Relative sensitivity of the estimated net reduction in atmospheric $CO_2$ to uncertainty in key sequestration and avoided emission parameters for each scenario. Sensitivity is assessed relative to a model run with no uncertainty, in which model parameters are set at their central estimate. A value of 1 indicates the estimated net reduction in $CO_2$ is the same as a model run without uncertainty, while a value of 0.5 or 2 would represent a halving or doubling of estimated climate benefits, respectively. Violin plots show the distribution of estimates from all Monte Carlo runs, with each individual estimate shown as a shaded point. Note the scale of the y-axis varies between panels.

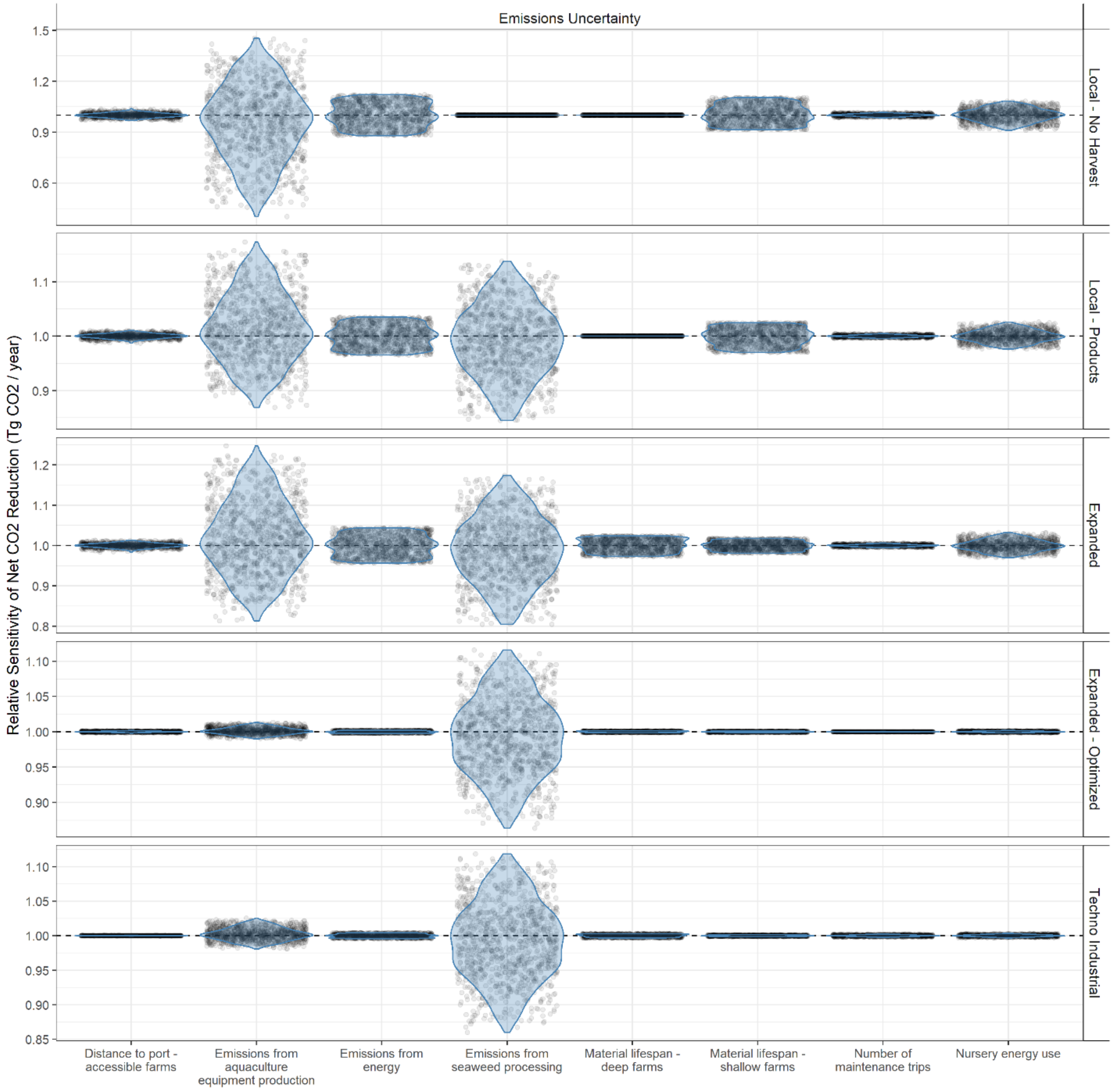


**Fig. S4C.**

Relative sensitivity of the estimated net reduction in atmospheric $CO_2$ to uncertainty in key emissions parameters for each scenario. Sensitivity is assessed relative to a model run with no uncertainty, in which model parameters are set at their central estimate. A value of 1 indicates the estimated net reduction in $CO_2$ is the same as a model run without uncertainty, while a value of 0.5 or 2 would represent a halving or doubling of estimated climate benefits, respectively. Violin plots show the distribution of estimates from all Monte Carlo runs, with each individual estimate shown as a shaded point. Note the scale of the y-axis varies between panels.

**Table S1.**

Summary values for key outputs from Monte Carlo runs of each scenario. Values represent the 25th, 50th (median), and 75th percentile of 10,000 Monte Carlo runs. These values correspond to the values reported for the *Expanded* scenario in Figure 2 of the main text.

| | | **Local - No Harvest** | | | **Local - Products** | | | **Expanded** | | | **Expanded - Optimized** | | | **Techno Industrial** | | |
|---|---|---|---|---|---|---|---|---|---|---|---|---|---|---|---|---|
| | Units | 25% | Median | 75% | 25% | Median | 75% | 25% | Median | 75% | 25% | Median | 75% | 25% | Median | 75% |
| Net Primary Productivity | Tg C / year | 0.0122 | 0.0214 | 0.0348 | 0.0122 | 0.0214 | 0.0348 | 0.0292 | 0.0511 | 0.0831 | 0.3986 | 0.5967 | 0.8706 | 1.8715 | 2.8014 | 4.0876 |
| Harvested Biomass | Tg ww / year | 0.2397 | 0.4060 | 0.6072 | 0.2397 | 0.4060 | 0.6072 | 0.5722 | 0.9689 | 1.4491 | 7.2779 | 10.7202 | 14.4696 | 34.1702 | 50.3318 | 67.9355 |
| DOC and POC | Tg C / year | 0.0051 | 0.0096 | 0.0172 | 0.0051 | 0.0096 | 0.0172 | 0.0121 | 0.0229 | 0.0411 | 0.1633 | 0.2685 | 0.4442 | 0.7666 | 1.2607 | 2.0854 |
| Passive sequestration | Tg $CO_2$ / year | 0.0021 | 0.0041 | 0.0076 | 0.0021 | 0.0041 | 0.0076 | 0.0057 | 0.0110 | 0.0202 | 0.0762 | 0.1296 | 0.2206 | 0.3656 | 0.6222 | 1.0595 |
| Deep sequestration | Tg $CO_2$ / year | - | - | - | - | - | - | - | - | - | - | - | - | 0.4368 | 0.7061 | 1.1008 |
| On-shelf sequestration | Tg $CO_2$ / year | 0.0026 | 0.0046 | 0.0076 | 0.0003 | 0.0005 | 0.0008 | 0.0007 | 0.0012 | 0.0020 | 0.0093 | 0.0143 | 0.0216 | - | - | - |
| Seaweed Products CO2 Avoided | Tg $CO_2$ / year | - | - | - | 0.0775 | 0.1358 | 0.2104 | 0.1679 | 0.2895 | 0.4420 | 2.2486 | 3.3366 | 4.6299 | 6.0287 | 8.9003 | 12.2701 |
| Total CO2 sequestered or avoided | Tg $CO_2$ / year | 0.0051 | 0.0093 | 0.0154 | 0.0813 | 0.1422 | 0.2196 | 0.1772 | 0.3062 | 0.4660 | 2.3817 | 3.5164 | 4.8731 | 6.5983 | 9.7051 | 13.3551 |
| Nursery Emissions | Tg $CO_2$ / year | 0.0031 | 0.0045 | 0.0060 | 0.0031 | 0.0045 | 0.0060 | 0.0074 | 0.0107 | 0.0144 | 0.0074 | 0.0108 | 0.0144 | 0.0350 | 0.0506 | 0.0675 |
| Cultivation Emissions | Tg $CO_2$ / year | 0.0184 | 0.0222 | 0.0259 | 0.0184 | 0.0222 | 0.0259 | 0.0495 | 0.0598 | 0.0694 | 0.0493 | 0.0597 | 0.0695 | 0.2644 | 0.3158 | 0.3652 |
| Transport and Processing Emissions | Tg $CO_2$ / year | - | - | - | 0.0070 | 0.0134 | 0.0223 | 0.0167 | 0.0320 | 0.0531 | 0.2148 | 0.3610 | 0.5536 | 0.6264 | 1.0297 | 1.5504 |
| Total emissions | Tg $CO_2$ / year | 0.0228 | 0.0267 | 0.0308 | 0.0334 | 0.0409 | 0.0503 | 0.0862 | 0.1045 | 0.1274 | 0.2853 | 0.4320 | 0.6241 | 0.9949 | 1.4001 | 1.9180 |
| Net reduction in CO2 | Tg $CO_2$ / year | -0.0225 | -0.0166 | -0.0096 | 0.0447 | 0.0995 | 0.1718 | 0.0840 | 0.1961 | 0.3447 | 2.0185 | 3.0396 | 4.2909 | 5.3925 | 8.1513 | 11.5315 |

**Table S2.**

Gross and net (accounting for emissions) estimated climate benefit for each possible seaweed fate per km$^2$ of cultivated area (T $CO_2$e / km$^2$ / year), assuming all seaweed biomass is directed towards a single fate. Results are shown for Expanded and Expanded-Optimized scenarios only, with the difference between the scenarios only in the kelp production rate. These scenarios are largely indicative of all other scenarios. Values are reported as the median estimate from 10,000 Monte Carlo runs with 25$^{th}$ and 75$^{th}$ quantiles in brackets. Negative values for net benefits indicate an overall release of atmospheric greenhouse gases.

| | Fate | Climate Benefits (T $CO_2$e / km$^2$ / year) | |
|---|---|---|---|
| | | **Expanded** | **Expanded - Optimized** |
| Gross | Food products | 315.9 (179.3-495.3) | 3665 (2416.3-5221.6) |
| | Animal Feeds | 242 (145.1-360) | 2787 (1940.5-3727.8) |
| | Fuels | 89.6 (53.4-133.5) | 1032.4 (721-1383.4) |
| | On-shelf release | 20.4 (11.3-33.9) | 240.3 (153.6-360.6) |
| | Deep release | 32.2 (17.7-53.6) | 375.9 (241.1-567.6) |
| Net (Gross - Emissions) | Food products | 223.9 (99.5-392.2) | 3237.7 (2084.8-4696.2) |
| | Animal Feeds | 150.2 (64.7-252.1) | 2350.2 (1619.1-3166.4) |
| | Fuels | -3.3 (-28.4-27.4) | 582.4 (370.4-852.6) |
| | On-shelf release | -36.1 (-48.8 - -20.8) | 181.8 (95.8-304) |
| | Deep release | -26.5 (-42.8 - -5.4) | 296.4 (164.4-482.8) |

**Table S3.**

Full list of model parameters. Column SD provides the standard deviation, where available. Column Dist indicates the shape of the distribution used for the parameter; triangular (tri), truncated normal (tnorm), or uniform (unif). The sources of the parameter values are also provided.

| Module | Parameter | Parameter Description | Value | Units | Min | Max | SD | Dist | Source |
|---|---|---|---|---|---|---|---|---|---|
| Production | $S_S$ | Pct area growing Sacch | 0.8 | NA | | | 0 | NA | Survey |
| Production | $S_A$ | Pct area growing Alaria | 0.1 | NA | | | 0 | NA | Survey |
| Production | $S_N$ | Pct area growing Nereo | 0.1 | NA | | | 0 | NA | Survey |
| Production | $H_n$ | Number of harvests per year | 1 | NA | | | 0 | NA | Survey |
| Production | $FL_{POC}$ | Fraction seaweed lost as POC | 0.167 | kg POC / kg C | 0 | 0.55 | | tri | [4] |
| Production | $FL_{DOC}$ | Fraction seaweed lost as DOC | 0.233 | kg DOC / kg C | 0.137 | 0.342 | | tri | [4] |
| Production | C | Carbon content | 0.248 | kg C / kg dw | | | 0.063 | tnorm | [3] |
| Production | $B_{A.S}$ | Area-based harvest Sacch | 0.78 | kg ww /m2 / yr | | | 0.801 | tnorm | Survey |
| Production | $B_{A.A}$ | Area-based harvest Alaria | 0.22 | kg ww /m2 / yr | | | 0.134 | tnorm | Survey |
| Production | $B_{A.N}$ | Area-based harvest Nereo | 0.26 | kg ww /m2 / yr | | | 0.26 | tnorm | Survey |
| Production | $B_{A.S}$ - Lit | Area-based harvest Sacch – literature | 8.3 | kg ww /m2 / yr | | | 5.9 | NA | Various (see Table S5) |
| Production | $B_{A.A}$ - Lit | Area-based harvest Alaria – literature | 8.3 | kg ww /m2 / yr | | | 5.9 | NA | Various (see Table S5) |
| Production | $B_{A.N}$ - Lit | Area-based harvest Nereo – literature | 8.3 | kg ww /m2 / yr | | | 5.9 | NA | Various (see Table S5) |
| Sequestration | $F_{DOC.Seq}$ | Fraction DOC sequestered | 0.14 | kg C / kg C | 0.106 | 0.402 | | tri | [6] |
| Sequestration | $F_{POC.Seq}$ | Fraction POC sequestered | 0.14 | kg C / kg C | 0.106 | 0.402 | | tri | [6] |

| Module | Parameter | Parameter Description | Value | Units | Min | Max | SD | Dist | Source |
|---|---|---|---|---|---|---|---|---|---|
| Sequestration | $F_{Rep.Food}$ | Food replacement factor | 3.5 | kg CO2 / kg DW | 1 | 6 | | tri | [10] |
| Sequestration | $F_{Rep.Feed}$ | Feed replacement factor | 2.5 | kg CO2 / kg DW | 2 | 3 | | tri | [10] |
| Sequestration | $F_{Rep.Fuel}$ | Fuel replacement factor | 0.85 | kg CO2 / kg DW | 0.7 | 1 | | tri | [10] |
| Sequestration | $K_C$ | Fraction seaweed carbon from atmospheric source | 0.5 | % | 0.4 | 0.8 | | tri | [10,11] |
| Sequestration | $K_{C.CO2}$ | Carbon to CO2 conversion | 3.67 | kg CO2 / kg C | | | | NA | NA |
| Emissions | $NRG_{Nurs}$ | Nursery daily energy use | 0.37 | GWh / km2 / year | 0.195 | 0.544 | | tri | [13,14] |
| Emissions | $E_{Nrg}$ | CO2 emissions per energy | 24900 | kg CO2e / GWh | 9700 | 40100 | | unif | [24] |
| Emissions | $E_{Mat}$ | CO2 emissions material production | 44070 | kg CO2e / km2 | 15990 | 65520 | | tri | [16,18,19] |
| Emissions | $M_{Mat}$ | Annual mass of material | 766.5 | kg / km2 / yr | 378 | 1155 | | tri | [16,20] |
| Emissions | $E_{Barge}$ | CO2 emissions barge transport | 3E-05 | kg CO2 / kg / km | 3E-05 | 3E-05 | | tri | [10,19] |
| Emissions | $E_{Ves}$ | CO2 emissions maintenance vessel | 5.0427 | kg CO2 / km | 2.3653 | 7.72 | | tri | [10,19] |
| Emissions | $N_{Maint}$ | Number maintenance trips | 6 | trips/year | 3 | 9 | | tri | [20] |
| Emissions | $A_{Maint}$ | Area maintained per trip | 0.5 | km2 | 0.25 | 0.75 | | tri | [20] |
| Emissions | $D_{Maint}$ | Distance traveled per km maintained | 10 | km/km2 | 5 | 15 | | tri | [20] |

| Module | Parameter | Parameter Description | Value | Units | Min | Max | SD | Dist | Source |
|---|---|---|---|---|---|---|---|---|---|
| Emissions | $E_{Sink}$ | CO2 emissions from sinking mechanism | 5E-05 | kg CO2 / kg ww | 0 | 0.000 1 | | tri | Calculated |
| Emissions | $E_{Conv}$ | CO2 emissions seaweed processing | 0.039 | kg CO2 / kg ww | 0.0009 | 0.082 7 | | tri | [10,14,23] |
| General | $DW_S$ | Wet to dry conversion ratio Sacch | 0.113 | kg dw / kg ww | | | | NA | [2] |
| General | $DW_A$ | Wet to dry conversion ratio Alaria | 0.165 | kg dw / kg ww | | | | NA | [1] |
| General | $DW_N$ | Wet to dry conversion ratio Nereo | 0.129 | kg dw / kg ww | | | | NA | [1] |
| Production | A - shallow and near | Area for shallow farms near to communities | 5,070 | km2 | | | | NA | Calculated |
| Production | A – deep and near | Area for deep farms near to communities | 7,030 | km2 | | | | NA | Calculated |
| Production | A – shallow and distant | Area for shallow farms distant to communities | 11,610 | km2 | | | | NA | Calculated |
| Production | A – deep and distant | Area for deep farms distant to communities | 33,100 | km2 | | | | NA | Calculated |
| Emissions | $D_{Port}$ - near | Distance from port for near farms | 12.5 | km | | | 5 | tnorm | Calculated |
| Emissions | $D_{Port}$ - distant | Distance from port for distant farms | 62.5 | km | | | 15 | tnorm | Calculated |
| Emissions | $D_{Sink}$ - shallow | Distance to sink location shallow | 100 | km | | | 35 | tnorm | Calculated |
| Emissions | $D_{Sin}$ - deep | Distance to sink location deep | 75 | km | | | 25 | tnorm | Calculated |
| Sequestration | Seq.Mult - shallow | Passive seq multiplier shallow farm | 1 | NA | 1 | 1 | | unif | Calculated |
| Sequestration | Seq.Mult - deep | Passive seq multiplier deep farm | 1.2 | NA | 1.1 | 1.3 | | unif | Calculated |
| Emissions | Mat.Life - shallow | Material depth multiplier shallow | 1 | NA | 0.9 | 1.1 | | unif | Calculated |
| Emissions | Mat.Life - deep | Material depth multiplier deep | 0.8 | NA | 0.7 | 0.9 | | unif | Calculated |
| Sequestration | $F_{Active.Seq}$ - deep | Fraction deep released seaweed sequestered >500 | 0.47 | kg C / kg C | 0.1 | 0.97 | | tri | [8] |
| Sequestration | $F_{Active.Seq}$ - shallow | Fraction shallow released seaweed sequestered | 0.14 | kg C / kg C | 0.106 | 0.402 | | tri | [6] |

**Table S4.**

Area-based biomass production rates (kg ww / $m^2$) from regional seaweed producers. For Nereocystis we received only one production estimate and therefore assumed the standard deviation was equivalent to the mean.

| Species | Number of Values | Mean | Standard Deviation |
|---|---|---|---|
| Saccharina | 10 | 0.78 | 0.801 |
| Alaria | 3 | 0.22 | 0.134 |
| Nereocystis * | 1 | 0.26 | 0.260 |

**Table S5.**

Area-based biomass production rates (kg ww / $m^2$) compiled from the literature.

| Species | Location of Study | Value (kg ww / $m^2$) | Source | Notes |
|---|---|---|---|---|
| Unspecified | Model-based | 20 | [25] | Estimated for Macrocystis from Chile |
| Generic 'Temperate Brown' model group | Model-based | 13.5 | [26] | Original estimate (1.35 kg DW/$m^2$) was in dry weight, converted to wet weight conservatively assuming a ratio of 1:10 |
| *Macrocystis pyrifera* | Chile | 12.4 | [27] | Maximum yield reported by authors |
| *Macrocystis pyrifera* | US West Coast | 10 | [7] | Original estimate (1 kg DW/$m^2$) was in dry weight, converted to wet weight conservatively assuming a ratio of 1:10 |
| *Saccharina latissima* | US East Coast | 8 | [28] | Original estimate (24 kg/m) was for single line, we assumed 3 m line spacing to calculate area based production rates |
| *Saccharina latissima* | Model-based | 7.2 | [16] | Original estimate (72 T/Ha) was for single layer, also estimated 108 T/Ha (10.8 kg /$m^2$) for double layer system. |
| *Saccharina latissima* | US East Coast | 4.2 | [29] | Original estimate (12.7 kg/m) was for single line, we assumed 3m line spacing to calculate areal production rates |
| Unspecified | Model-based | 4.2 | [19] | Original estimate (12.5 kg/m) was for single line, we assumed 3m line spacing to calculate areal production rates |
| *Saccharina latissima* | Alaska | 1.7 | [30] | Original estimate (5 kg/m) was for single line, we assumed 3m line spacing to calculate areal production rates |
| *Saccharina latissimi & Alaria esculenta* | Faroe Islands | 1.4 | [31] | Original estimate (1437.5 kg DW/Ha) was in dry weight, converted to wet weight conservatively assuming a ratio of 1:10 |
| **Mean** | | 8.3 | | |
| **Standard Deviation** | | 5.9 | | |

**Table S6.**

Questions included in the questionnaire sent to kelp producers in British Columbia, Canada and Alaska, USA.

| | |
|---|---|
| 1) | Where is your farm located (geographic coordinates preferred)? If you operate multiple farms, please list them all, or complete a questionnaire for each farm. |
| 2) | What species do you farm? What is the spatial footprint (marine area) for each species you farm? |
| 3) | How much do you produce annually for each species, either in total (e.g., kg) , or by line density (e.g., kg/m of line)? |
| 4) | When do you typically harvest each species? What informs your decision of when to harvest? Can you do multiple harvests per year? Why or why not? |
| 5) | Please describe your farm configuration (by species if needed), including number of horizontal lines, line spacing, and line length. |
| 6) | Do you use vertical lines or any other non-standard farm configuration? Please provide as much detail as you are able. |
| 7) | Can you describe your seeding and out-planting operation? Do you use direct or spore seeding? What are the main trade-offs between these approaches for your operation? |
| 8) | We estimate small farms would be < 50 acres, mid-size farms up to 100 acres, and large, potentially offshore farms exceeding (1 $km^2$, or 250 acres). Do you agree with these classes? Would you distinguish between types of farms in some other way? Provide as much information as you like. |
| 9) | How are bigger farms different from smaller farms - specifically in terms of seaweed production? Please provide as much information as you can. |
| 10) | Value-added seaweed (e.g., food, biofuel, bioplastics, animal feed, fertilizer, biochar) sold to markets as an alternative to existing, more carbon-intense products. Please provide as much detail as you can on the products and the anticipated net carbon reduction. |
| 11) | Seaweeds transported to a specific ocean location at a specified depth for potential carbon credits. |
| 12) | Seaweeds transported to a specific ocean location at the surface for potential carbon credits. |
| 13) | Seaweed released near the farm for potential carbon credits. |
| 14) | If you are planning to sequester carbon in the ocean, how far are potential carbon sequestration sites from your farm(s)? Please provide as much detail as you can. |
| 15) | How do you power your seaweed nursery facilities? Can you estimate annual energy use? |
| 16) | How much fuel (in volume or dollars) did your vessels use last season (total across all vessels including on-farm operations, maintenance, and transport of harvest to the dock or to a processor)? |
| 17) | Can you estimate the proportion of this vessel fuel used for 1) operations (besides nursery facilities), 2) maintenance, and 3) transport of harvest? |

18) If you process your harvest at your own facility, do you use:

19) If you process your harvest at your own facility, how much electricity from the grid did you use for processing last year? (kWh or dollars)? How much diesel (volume or dollars) did you use for generators? Did you use any other sources of energy?

20) Do you transport unprocessed seaweed to a processor by road? If so, how long is the trip?

21) Can you estimate your annual capital equipment replacement costs (i.e., equipment might include lines, buoys, boats, docks, etc.)? Please provide as much detail as you are able.

22) How is the farm maintained during production? Can you estimate the energy used for one maintenance trip? What area of the farm can you maintain during each trip? How many such trips (on average) are required for one harvest?

23) How might bigger farms different from smaller farms - specifically in terms of operating costs? Please provide as much information as you can.

24) Please provide any additional comments you think may be relevant to this modelling effort.
